# Matriptase generates a tissue damage response via promoting Gq signalling, leading to RSK and DUOX activation

**DOI:** 10.1101/2021.01.21.427549

**Authors:** MA Jiajia, Claire A. Scott, HO Ying Na, Harsha Mahabaleshwar, Katherine S. Marsay, Changqing Zhang, Christopher K. J. Teow, NG Ser Sue, Weibin Zhang, Vinay Tergaonkar, Lynda J. Partridge, Sudipto Roy, Enrique Amaya, Tom J. Carney

## Abstract

Tissues respond to damage by increasing inflammation and epithelial cell motility. How damage detection and responses are orchestrated is unclear. Overexpression of the membrane bound protease, Matriptase, or mutation of its inhibitor, Hai1, results in inflamed epithelia, in which cells have increased motility and are prone to carcinoma. How Matriptase leads to these cellular outcomes is unknown. We demonstrate that zebrafish *hai1a* mutants show increased H_2_O_2_, NfκB signalling, and IP_3_R-mediated calcium flashes, and that these promote inflammation, but do not generate epithelial cell motility. In contrast, inhibition of the Gq subunit rescues both the *hai1a* inflammation and epithelial phenotypes, with the latter recapitulated by the DAG analogue, PMA. We demonstrate that *hai1a* has elevated pERK, inhibition of which rescues the epidermal defects. Finally, we identify RSK kinases as pERK targets disrupting adherens junctions in *hai1a* mutants. Our work maps novel signalling cascades mediating the potent effects of Matriptase on epithelia, with implications for tissue damage response and carcinoma progression.

## Introduction

Cells express diverse systems on their cell surface that ensure they can detect and respond to local tissue damage. Dysregulation of these systems are known to promote tumour development, and tumours have been proposed to represent non-healing wounds. Type II transmembrane serine proteases have been implicated in many diverse physiological processes and pathologies, including iron homeostasis, epithelial barrier formation, hypertension, cancer, and viral infection (Szabo & Bugge, 2008). Of these, Matriptase, encoded by the *ST14* gene, has been extensively studied due to its broad expression in most epithelia, its consistent dysregulation in human carcinomas and its ability to promote malignancy when overexpressed in mice (List et al., 2005; Martin & List, 2019). Matriptase is essential for epidermal cornification and desquamation, with *ST14* mutations found in patients with various forms of ichthyosis (Basel-Vanagaite et al., 2007). The oncogenic potency of Matriptase is mitigated by a Kunitz type transmembrane serine protease inhibitor, Hai1, encoded by the *SPINT1* gene. Indeed, loss of mouse Hai1 creates epidermal and intestinal barrier defects, and failure of placental labyrinth formation, which is due unrestricted Matriptase activity (Kawaguchi et al., 2011; Nagaike et al., 2008; Szabo, Molinolo, List, & Bugge, 2007). Mutation of the zebrafish orthologue Hai1a (encoded by *spint1a* gene) also results in epidermal defects, including removal of E-cadherin from the cell membrane leading to aberrant mesenchymal behaviour of keratinocytes, entosis, cell extrusion and sterile inflammation, which can all be ameliorated by reduction of Matriptase levels (Armistead, Hatzold, van Roye, Fahle, & Hammerschmidt, 2020; Carney et al., 2007; Mathias et al., 2007; Schepis et al., 2018). Similarly, overexpression of Matriptase in the mouse epidermis leads to epidermal papillomas, ulcerative and invasive carcinomas, and inflammation, suggesting there is a conserved response to unregulated Matriptase activity in keratinocytes of vertebrates (List et al., 2005). Clinically, an increase in the Matriptase:HAI-1 ratio has been found in a range of tumours, and is predictive of poor outcome (Martin & List, 2019).

Due to the clinical implications of its dysregulation, the targets of Matriptase, and the signalling pathways activated pathologically, are increasingly investigated. In transgenic mouse models and breast cancer cell lines, maturation of HGF by Matriptase has been implicated in activating a cMET-Gab1-PI3K-Akt-mTor pathway, resulting in carcinoma progression (Szabo et al., 2011; Zoratti et al., 2015). In contrast, cMet is not required for the effects of uninhibited Matriptase in zebrafish (Carney et al., 2007). In addition to HGF, elegant genetic analysis using mouse and zebrafish have established that the G-protein coupled receptor, Proteinase-activated receptor-2 (Par2; encoded by the *F2rl1* gene) is essential for the oncogenic and also the inflammatory effects of uninhibited Matriptase (Sales et al., 2015; Schepis et al., 2018). Like HGF, Par2 is a direct proteolytic target of Matriptase, which activates Par2 by cleaving within the extracellular N-terminal domain, removing an inhibitory peptide, and revealing a tethered ligand. Ligand bound Par2 then activates number of Gα protein subunits. It displays biased agonism, with different proteases/cleavage sites within the N-terminal domain activating distinct Gα subunits (Pawar, Buzza, & Antalis, 2019). Early studies in keratinocytes linked Par2 activation with intracellular Ca^++^ mobilization via phospholipase C, thus implicating Gq subunit as the canonical target (Schechter, Brass, Lavker, & Jensen, 1998). Alternate Gα subunits, including Gi. Gs and G12/13 are now known to also be activated by Par2, thus placing cAMP and Rho pathways downstream of Par2 (Zhao, Metcalf, & Bunnett, 2014). The logic of the pathway utilised depends on cell context and the activating protease. *In vitro* analyses implicated both Par2 and Gi in Matriptase mediated Nfκb pathway activation. This work however excluded involvement of other Gα subunits *in vitro* (Sales et al., 2015). Whilst this explains the inflammatory phenotype of uninhibited Matritase1, it fails to explain how Par2 suppresses the carcinoma phenotypes in the epidermis. It is also unclear how Par2 and HGF both contribute to the pro-oncogenic effects of Matriptase. In addition to Gα subunits, Par2 can transactivate EGFR through an unknown mechanism, and inhibition of EGFR alleviates certain basal keratinocyte phenotypes of zebrafish *hai1a* mutants (Schepis et al., 2018). Thus, the identity, contribution, and interactions of the pathways downstream of Matriptase remain unclear.

We have utilized the zebrafish *hai1a* mutant as an *in vivo* platform to determine mechanisms and pathways activated by Matriptase. Informed by an unbiased proteomics approach, we demonstrate that Gq is essential for all phenotypes, with IP_3_R dependent Ca^++^ release predominantly accounting for DUOX and Nfκb mediated sterile inflammation, whilst PKC and pERK drive the epithelial defects. We demonstrate that inhibition of RSK kinases, known pERK targets, also rescues the epithelial defects and restore E-cadherin to the membrane of basal keratinocytes in *hai1a* mutants. This has allowed us to construct a framework of the pathways downstream of zebrafish Matriptase.

## Results

### Increased hydrogen peroxide and calcium flashes contribute to inflammation in *hai1a* mutants

The *hai1a* zebrafish mutant combines epidermal defects and inflammation through loss of Matriptase1a inhibition, and subsequent activation of the GPCR, Par2b. Using the *Tg(mpx:eGFP)*^*i114*^ transgenic line to label neutrophils, we examined the inflammation phenotype to determine the mechanistic basis. As previously shown, neutrophils in *hai1a* occupy the epidermis, are highly active and motile, but move randomly (Mathias et al., 2007) (Fig. 1A-E; Video 1). Neutrophils in *hai1a* are successfully recruited locally to a small needle wound of the larval fin but were not recruited from further away or from the vasculature as in wild-type (Mathias et al., 2007). To better compare the signals recruiting neutrophils to a wound and those activating neutrophils in *hai1a* we performed a larger fin wound in wild-type and *hai1a* mutants by scalpel cut of the tail fin at 4dpf. We noted that again, whilst there was brief local recruitment in the mutant, more distant neutrophils appeared apathetic to the wound and remained migrating randomly, unlike in wild-type where neutrophils were recruited from a great distance and tracked to the wound with high directionality (Fig. 1F, G; Video 2). We noted, however, that although *hai1a* mutant neutrophils lacked directionality following wounding, they did increase their speed during their random movements, suggesting other damage response components present (Fig. S1A). We further tested if the early recruitment of neutrophils to the epidermis in *hai1a* mutants was dependent on apoptosis by performing a triple staining for neutrophils labelled by *Tg(fli1:EGFP)*^*y1*^ with an immunofluorescent antibody labelling against the basal keratinocyte transcription factor TP63, and TUNEL labelling of apoptotic cells. Whilst the epidermis of *hai1a* mutants, unlike WT, had regions of apoptosis at 24hpf (arrowhead, Fig. 1H, I), neutrophils were not associated, but rather found at nascent TUNEL negative aggregates of basal keratinocytes (arrow). We conclude that epidermal cell death does not directly contribute to inflammation and that the effector stimulating neutrophils in *hai1a* mutants is as, or more, potent as that of wounds.

**Figure 1:**
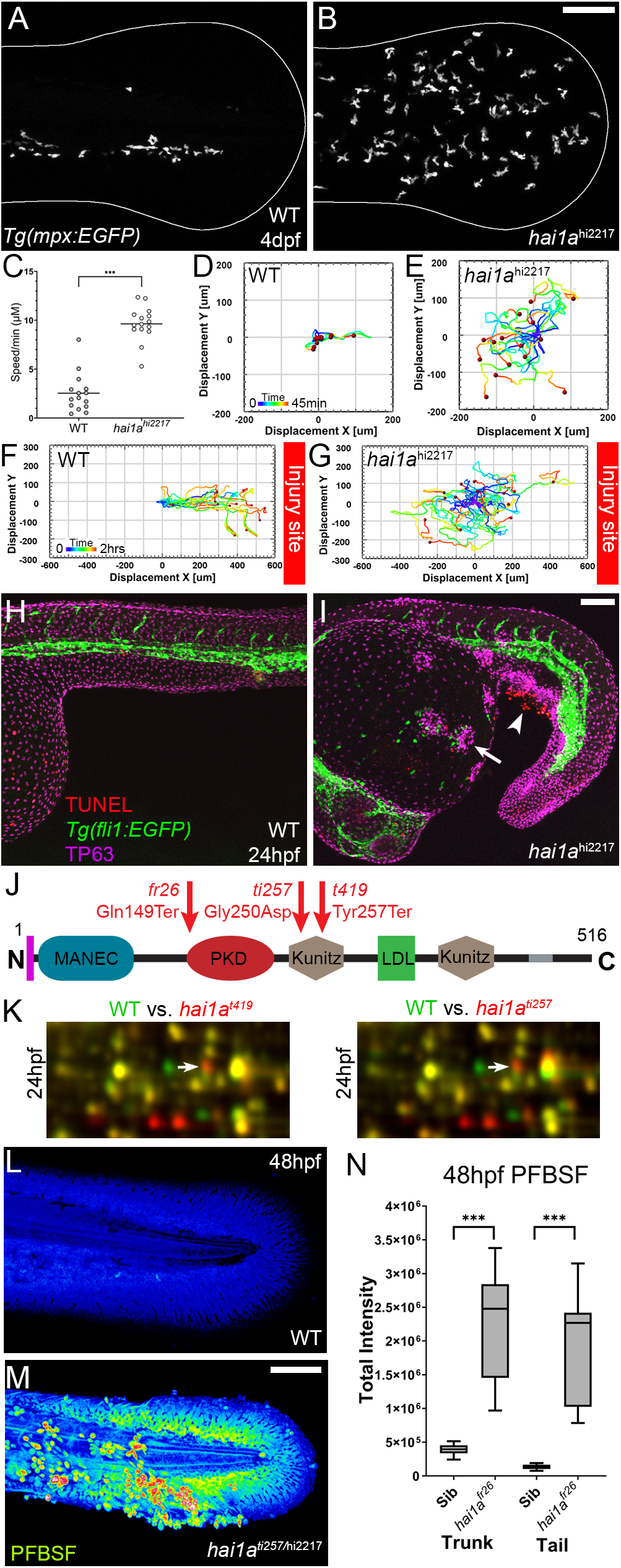
The epidermis of *hai1a* mutants displays elevated hydrogen peroxide. **A-B:** Projected confocal images showing neutrophils populate the tail of *hai1a*^*hi2217*^ mutants (B) but just the vasculature of WT (A) at 4dpf labelled by the *Tg(mpx:EGFP)*^*i114*^ line. Fin extremity outlined in white. **C:** Neutrophils move significantly faster in *hai1a*^*hi2217*^ than WT. n=15; t-test; *** = p<0.001. **D-E:** Tracks of neutrophil migration taken from Video 1 in WT (D) and *hai1a*^*hi2217*^ (E). **F-G:** Tracks of neutrophil migration taken from Video 2 in WT (F) and *hai1a*^*hi2217*^ (G) from immediately after wounding tail fin at indicated site. **H-I:** Projected confocal images showing neutrophils (green, labelled by *Tg(fli1:egfp)*^*y1*^ transgenic line) are not attracted to regions of apoptosis (arrowhead, red, stained by TUNEL), but to epidermal aggregates (arrow, magenta, stained by anti-TP63) in 24hpf *hai1a*^*hi2217*^ mutants (I) but not WT (H). **J**: Schematic of the Hai1a protein with protein domains given, signal peptide as purple line and trans-membrane domain as grey line. Location and nature of the *fr26* and two *dandruff* alleles, *ti257* and *t419* given. **K:** Selected region of 2D gel of protein extracted from 24hpf embryos for *hai1a*^*t419*^ (left) or *hai1a*^*ti257*^ (right) in red, superimposed over WT protein samples (green in both). The shift in pI of Peroxiredoxin4 in both alleles is indicated with an arrow. **L-M:** Projected lateral confocal views of PFBSF staining of WT (L) and *hai1ati257/hi2217* (M) tail fins at 48hpf. **N:** Box and whiskers plot of PFBSF fluorescent staining intensity of WT and *hai1a*^*fr26*^ mutants at 48hpf in trunk and tail. n=9; t-test *** = p<0.001. Scale bars B, I, M = 100μm

To identify the neutrophil activator in *hai1a* and visualize proteins cleaved by Matriptase1a, we employed an unbiased approach using 2D-gel proteomics to compare the wild-type proteome with that of strong *hai1a* alleles. The *dandruff (ddf)* mutant, isolated from an ENU screen, had many phenotypic similarities to the strong *hai1a*^*fr26*^ allele (van Eeden et al., 1996). Crosses between *ddf*^*ti257*^ or *ddf*^*t419*^ and *hai1a*^*hi2217*^ failed to compliment. Sequencing of *hai1a* cDNA from both *ddf* alleles identified a missense mutation (c.749G>A; p.Gly250Asp) in *ddf*^*ti257*^ and a nonsense mutation (c.771T>G; p.Tyr257Ter) in *ddf*^*t419*^ (Fig. 1J; Fig. S1B, C). The amino acid substituted in *ddf*^*ti257*^ is broadly conserved in Hai1 proteins of vertebrates suggesting its mutation is deleterious to function (Fig. S1D). We used both alleles for comparative 2D protein gel analysis at 24hpf and 48hpf. Rather than finding proteins with clear altered molecular weight, Peroxiredoxin4 (Prdx4) was identified as having a higher pI in both *hai1a*^*t419*^ and *hai1a*^*ti257*^ mutants at 24hpf and 48hpf, indicative of a change in oxidation state (Fig. 1F, Fig. S1E). Peroxiredoxins are hydrogen peroxide scavengers, and its altered oxidation state suggests that *hai1a* has higher H_2_O_2_ levels, a known activator of inflammation in larval zebrafish (Niethammer, Grabher, Look, & Mitchison, 2009). Using the stain pentafluorobenzenesulfonyl fluorescein (PFBSF) (Maeda et al., 2004), we observed significantly higher levels of H_2_O_2_ in the trunk and tails of *hai1a* mutants at 24 and 48hpf (Fig. 1G-I, Fig. S1G-H).

To demonstrate that, as with other phenotypes, this H_2_O_2_ increase in *hai1a* was due to unrestrained activity of Matriptase1a, we generated a *st14a (matriptase1a)* mutant by zinc finger mediated mutagenesis, which introduced 5bp into exon 6, altering the reading frame, and introducing a premature termination at 156 amino acids (*st14a*^*sq43*^; c.465_466insTCACA; p.Ala156SerfsTer8) (Fig. 2A, Fig. S2A-C). Homozygous zygotic *st14a* mutants showed no overt phenotype, however maternal zygotic mutants lacked ear otoliths (Fig. 2B-C). When crossed into the *hai1a* background, embryos lacking otoliths never displayed the *hai1a* epidermal phenotype (Table S1). Further, all *st14a*^*sq43*^; *hai1a*^*hi2217*^ double mutants lacked the epidermal and neutrophil phenotypes of *hai1a*^*hi2217*^ single mutants as expected (Fig. 2D-F). Double mutants also had significantly reduced H_2_O_2_ levels (Fig. 2F; Fig. S2D). To determine if this could account for the rescue of *hai1a* phenotypes, we used genetic and pharmacological inhibition of the main enzyme responsible for generating H_2_O_2_ in zebrafish, Duox. A morpholino directed against *duox* successfully reduced H_2_O_2_ levels (Fig. 2F; Fig. S2D) and neutrophil inflammation in *hai1a* mutants but did not rescue the epithelial defects (Fig. 2F-G). Treatment with a known Duox inhibitor, diphenyleneiodonium (DPI), also resulted in amelioration of neutrophil inflammation, but not epithelial aggregates in *hai1a* mutants (Fig. 2G, Fig. S2E). We conclude that excess H_2_O_2_ in *hai1a* mutants partially accounts for the neutrophil inflammation, but not epidermal defects.

**Figure 2:**
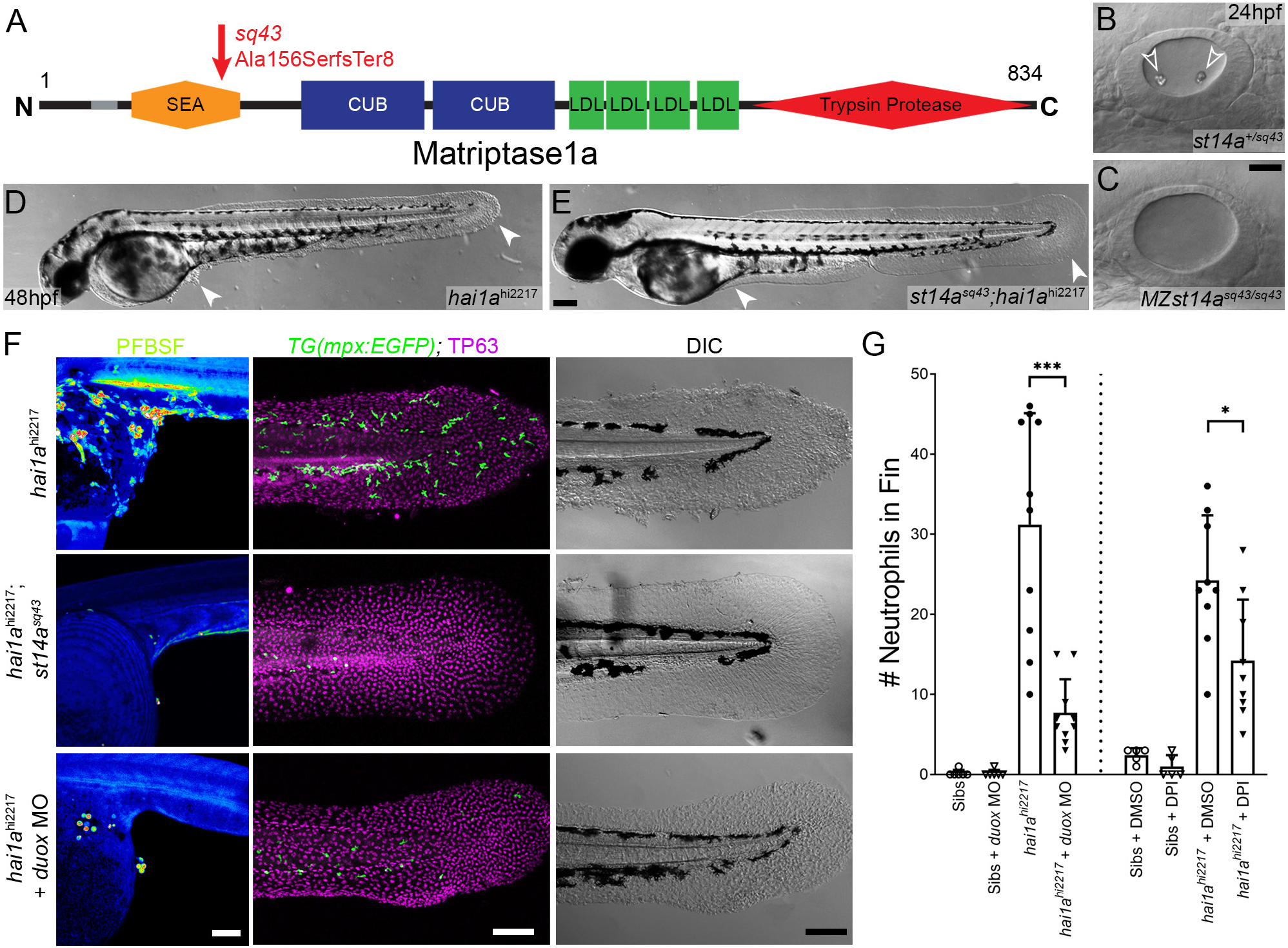
Loss of Matriptase1a or Duox1 reduces H_2_O_2_ and neutrophils in *hai1a* mutants. **A:** Schematic of the Matriptase1a protein with domains given and transmembrane domain as grey line. Location and nature of the *sq43* allele given by red arrow. **B-C:** Lateral DIC images of *st14a*^*+/sq43*^ (B) and MZ *st14a*^*sq43*^ (C) otic vesicles at 24hpf showing absence of otoliths (arrowheads in B) in the maternal zygotic *st14a* mutants. **D-E:** Lateral DIC images of *hai1a*^*hi2217*^ single mutant (D) and *st14a*^*sq43*^; *hai1a*^*h12217*^ double mutant (E) at 48hpf highlighting rescue of epidermal aggregates and fin morphology (arrowheads) in the double mutants. **F:** Projected confocal images of PFBSF staining at 24hpf (left column), TP63 (magenta) and eGFP (green) antibody staining at 48hpf (middle column) with DIC imaging (right column) for *hai1a*^*hi2217*^ single mutants (top row), *st14a*^*sq43*^; *hai1a*^*hi2217*^ double mutants (middle row) and *hai1a*^*hi2217*^ mutants injected with 0.4mM, *duox* MO + 0.2mM *tp53* morpholino (bottom row). Individuals for middle and right columns were hemizygous for the *Tg(mpx:EGFP)*^*i114*^ transgene. **G:** Counts of eGFP positive neutrophils on the fins of *hai1a*^*hi2217*^; *Tg(mpx:EGFP)*^*i114*^ or *Tg(mpx:EGFP)*^*i114*^, and either uninjected, injected with morpholino against *duox* (left side of graph), treated with 0.5% DMSO or 40μM DPI (right side of graph). n=10; t-test; *** = p<0.001; * = p<0.05. Scale bars C = 20μm; E, F = 100μm.

Duox is regulated by calcium through its EF-Hand domains, and calcium flashes are known to generate H_2_O_2_ in epidermal wounds in *Drosophila* (Razzell, Evans, Martin, & Wood, 2013). We injected *hai1a*^*fr26*^ with RNA encoding the calcium reporter *GCaMP6s*. Timelapse imaging at 24hpf indicated *hai1a* mutants had significantly more calcium flashes in both the trunk and tail (Fig. 3A, B, E; Video 3). Release of calcium from intracellular stores is regulated by IP3 receptors located on the endoplasmic reticulum. The frequency and number of calcium flashes in the trunk and tail of *hai1a* mutants is reduced by treatment with the IP_3_R inhibitor 2-APB compared to control (Fig. 3C, D, F; Video 4). Reducing calcium flashes in *hai1a* mutant embryos with 2-APB, also significantly reduced H_2_O_2_ levels (Fig. 3G; Fig. S3A), and partially reduced inflammation, however the epidermal defects were not noticeably rescued (imaged by DIC or labelled with the TP63 antibody) (Fig. 3G-I). We observed similar reduction in neutrophil inflammation, but not rescue of epidermal defects, in *hai1a* mutants treated with Thapsigargin, which inhibits the replenishment of ER calcium stores by SERCA (Fig. 3I; Fig. S3B-C). Thus, in *hai1a* mutants, IP_3_R dependent calcium flashes activate Duox, flooding the epidermis with H_2_O_2_ and leading to inflammation.

**Figure 3:**
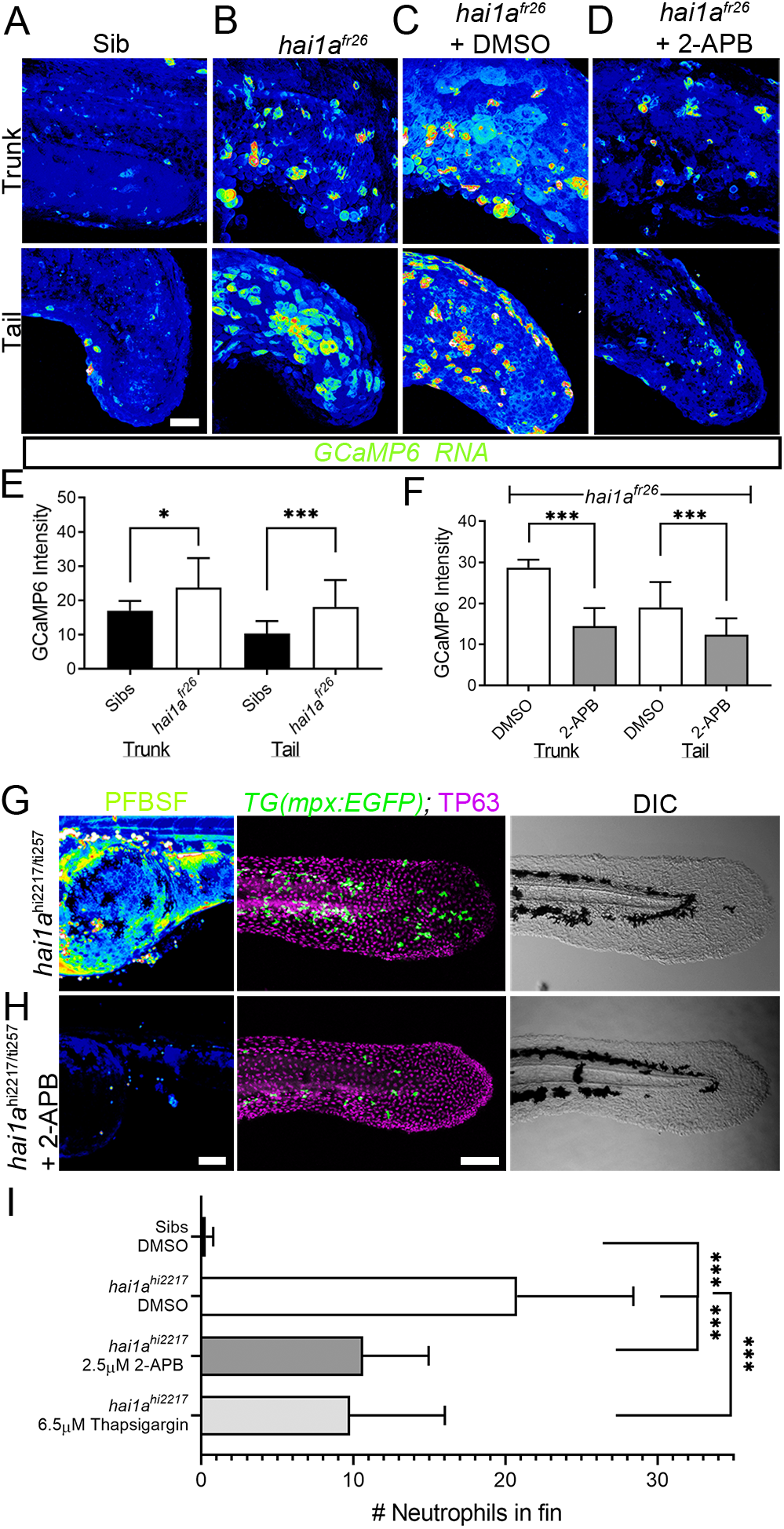
Calcium dynamics in *hai1a* mutants regulate H_2_O_2_ and inflammation. **A-D:** Projected confocal images of eGFP in the trunk (top row) or tail (bottom row) of WT (A), or *hai1a*^*fr26*^; (B-D) injected with *GCaMP6s* RNA, imaged at 24hpf indicating calcium dynamics. Embryos are either untreated (A-B), treated with DMSO (C), or with 2.5μM 2-APB (D). Images are temporal projections of timelapse movies taken at maximum speed intervals (2 minutes for tail movies, 3 minutes for trunk) and projected by time. **E-F:** Graphs comparing eGFP intensities from *GCaMP6s* RNA timelapses in trunk and tail between 24hpf WT and *hai1a*^*fr26*^ (E) and between *hai1a*^*fr26*^ treatedwith DMSO and 2.5μM 2-APB (F). n= 10; t-test; * = p<0.05, *** = p<0.001. **G-H:** Projected confocal images of PFBSF staining at 24hpf (left column), TP63 (magenta) and eGFP (green) antibody staining at 48hpf (middle column) with DIC imaging (right column) for *hai1a*^*hi2217/ti257*^ mutants (G), *or hai1a*^*hi2217/ti247*^ mutants treated with 2.5μM 2-APB (H). Individuals for middle and right columns were hemizygous for the *Tg(mpx:EGFP)*^*i114*^ transgene. **I:** Counts of eGFP positive neutrophils in the fins at 48hpf of *Tg(mpx:EGFP)*^*i114*^, or *hai1a*^*hi2217*^; *Tg(mpx:EGFP)*^*i114*^ treated with 0.5% DMSO, 2.5μM 2-APB or 6.5μM Thapsigargin. n=20; t-test; *** = p<0.001. Scale bars A-D = 50μm; G-H = 100μm.

### Hydrogen peroxide elevates NfkB signalling in hai1a mutants

Increased Matriptase, Par2 activity or hydrogen peroxide levels are known to activate NfkB signalling (Kanke et al., 2001; Sales et al., 2015; Schreck, Rieber, & Baeuerle, 1991). We crossed the *hai1a*^*fr26*^ allele to the NfkB sensor transgenic line *Tg(6xHsa.NFKB:EGFP)*^*nc1*^. In WT embryos, NfkB signalling was mostly restricted to neuromasts at 48hpf, whilst in *hai1a* mutants we observed an increase in fluorescence at 24hpf and a strong increase at 48hpf. Fluorescence at both timepoints was noted in epidermal aggregates and fin folds, locations of strong inflammation (Fig. 4A, B; Fig. S4A, B). This increase in signalling in 48hpf *hai1a* mutant embryos was confirmed by qRT-PCR of the NfkB target gene, *nfkbiaa* (Fig. 4C). To determine the extent that NfkB signalling accounts for the *hai1a* phenotypes, we generated a mutant in the *ikbkg* (= *ikkg* or *nemo*) gene by TALEN mediated mutagenesis, which encodes a scaffold protein required for activating the NfkB pathway (Rothwarf, Zandi, Natoli, & Karin, 1998) (*ikbkg*^*sq304*^ Gly80ValfsTer11; Fig. S4C). Crossing this mutant to *hai1a*^*hi2217*^ on the *Tg(mpx:eGFP)*^*i114*^ background realised a very strong rescue of neutrophil inflammation, but no improvement of *hai1a* epidermal defects (Fig. 4D-I). To demonstrate that this increase in NfkB signalling was dependent on H_2_O_2_, we injected *hai1a*^*hi2217*^; *Tg(6xHsa.NFKB:EGFP)*^*nc1*^ embryos with *duox* MO. We noted a strong reduction in NfkB pathway activation compared to uninjected *hai1a*^*hi2217*^ mutant controls (Fig. 4J, K). Conversely, genetic ablation of NfkB signalling, through use of the *ikbkg* mutant, did not reduce H_2_O_2_ levels in *hai1a* mutants (Fig. S4D-E). Similarly, we tested if reduction of calcium flashes could also reduce NfkB signalling in *hai1a* mutants using 2-APB, but noticed only a slight reduction (Fig. S4F-G). We conclude that upon loss of Hai1a, IP_3_R mediated release of calcium activates Duox to increase H_2_O_2_. This acts upstream of NfkB pathway activation, which is necessary for the inflammation phenotype.

**Figure 4:**
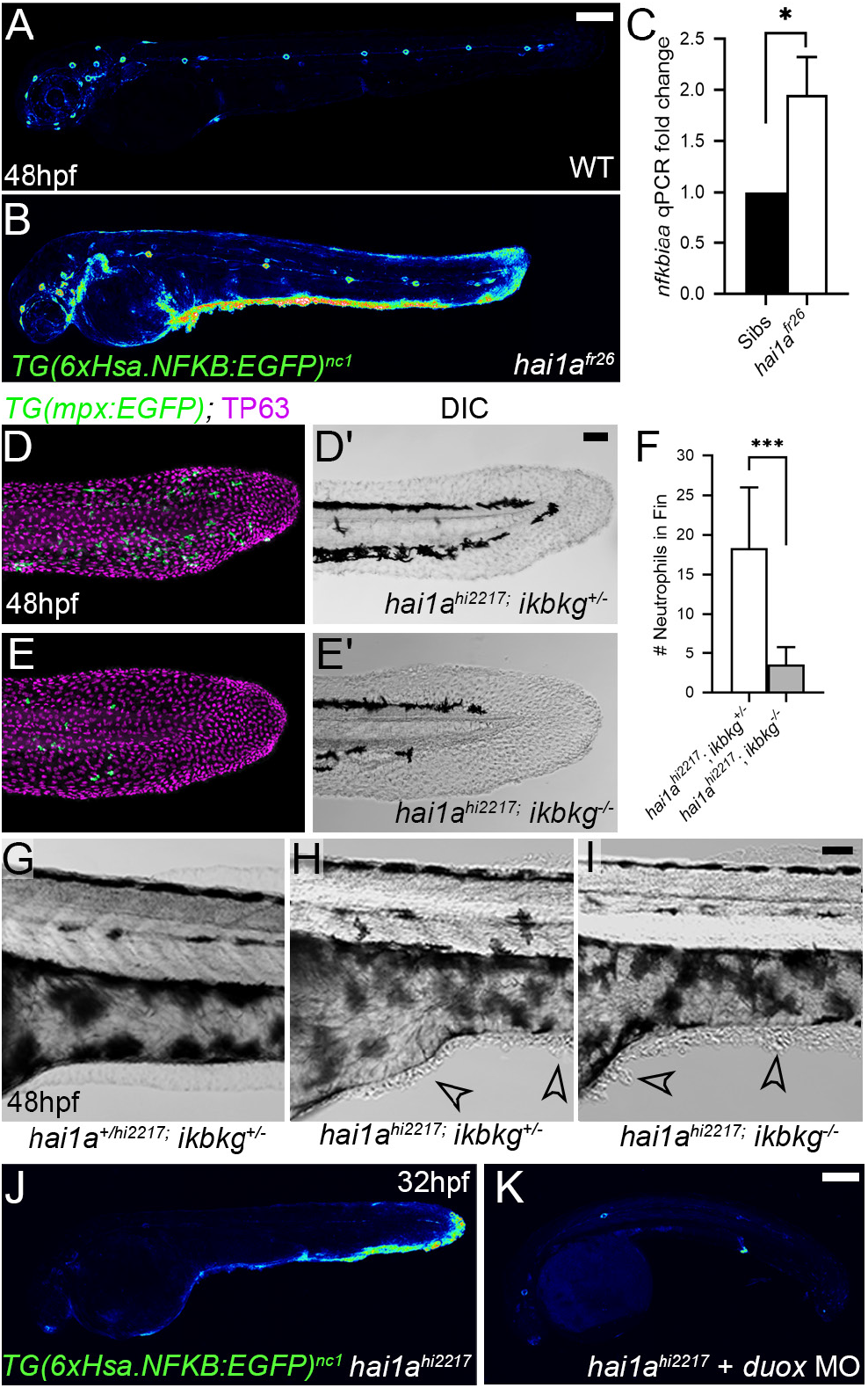
NfkB signalling is elevated in *hai1a* mutants and is necessary for neutrophil inflammation. **A-B:** Lateral confocal projections of *Tg(6xHsa.NFKB:EGFP)nc1* embryos reporting NfkB signalling levels at 48hpf for WT (A), and *hai1a*^*fr26*^ (B). **C:** qPCR of cDNA levels of NfkB target gene *nfkbiaa* in *hai1a*^*fr26*^ vs sibs at 48hpf. n=3, t-test *=p<0.05. **D-E’:** Projected confocal images of the tail fins of 48hpf *Tg(mpx:EGFP)*^*i114*^; *hai1a-hi2217* embryos, immunostained for TP63 (magenta) and eGFP (green) (D, E) with corresponding DIC image (D’, E’). Embryos were either mutant for *ikbkg (ikbkg*^*sq304*^, E-E’) or heterozygous (*ikbkg*^*+/sq304*^; D-D’). **F:** Counts of eGFP positive neutrophils in the fins at 48hpf of *hai1a*^*hi2217*^; *ikbkg*^*+/sq304*^ and *hai1a*^*hi2217*^; *ikbkg*^*sq304*^. Embryos were hemizygous for *Tg(mpx:EGFP)*^*i114*^. n=9; t-test; *** = p<0.001. **G-I:** Lateral DIC images of the trunk of *hai1a*^*+/hi2217*^; *ikb-kg*^*+/sq304*^ (G), *hai1a*^*hi2217*^; *ikbkg*^*+/sq304*^ (H), and *hai1a*^*hi2217*^; *ikbkg*^*sq304*^ (I). Loss of IKBKG does not rescue epidermal defects of *hai1a* mutants (arrowheads). **J-K:** Lateral confocal projections of *Tg(6xHsa. NFKB:EGFP)*^*nc1*^ embryos reporting NfkB signalling levels at 32hpf of *hai1a*^*hi2217*^ uninjected (J) or injected with *duox* MO (K). Loss of H_2_O_2_ reduces NfkB signalling levels in *hai1a* mutants. Scale bars A, K = 200μm; D’, I = 50μm.

### Both inflammation and epidermal aggregates of hai1a mutants are resolved by Gq inhibition

IP_3_ is generated from cleavage of PIP2 by Phospholipase C. The sensitivity of the *hai1a* mutants to 2-APB implies that IP_3_ levels are increased and therefore there may be an increase in Phospholipase C activity. Numerous attempts to inhibit PLC in *hai1a* mutants failed, and we were unable to find a dosage window that rescued without gross embryo deformity. Hence, we tested rescue of *hai1a* mutants with YM-254890, an inhibitor of the heterotrimeric G protein alpha subunit, Gq, which directly activates PLC isoforms. We found that not only did this significantly reduce neutrophil inflammation (Fig. 5D, F), but surprisingly, it also significantly rescued the epidermal defects in *hai1a* mutants (Fig. 5A-E). There was a visible reduction in epidermal aggregates in the trunk and improved tail fin fold integrity at 48hpf, either imaged by DIC or through TP63 immunostaining of basal keratinocytes (Fig. 5A-D).

**Figure 5:**
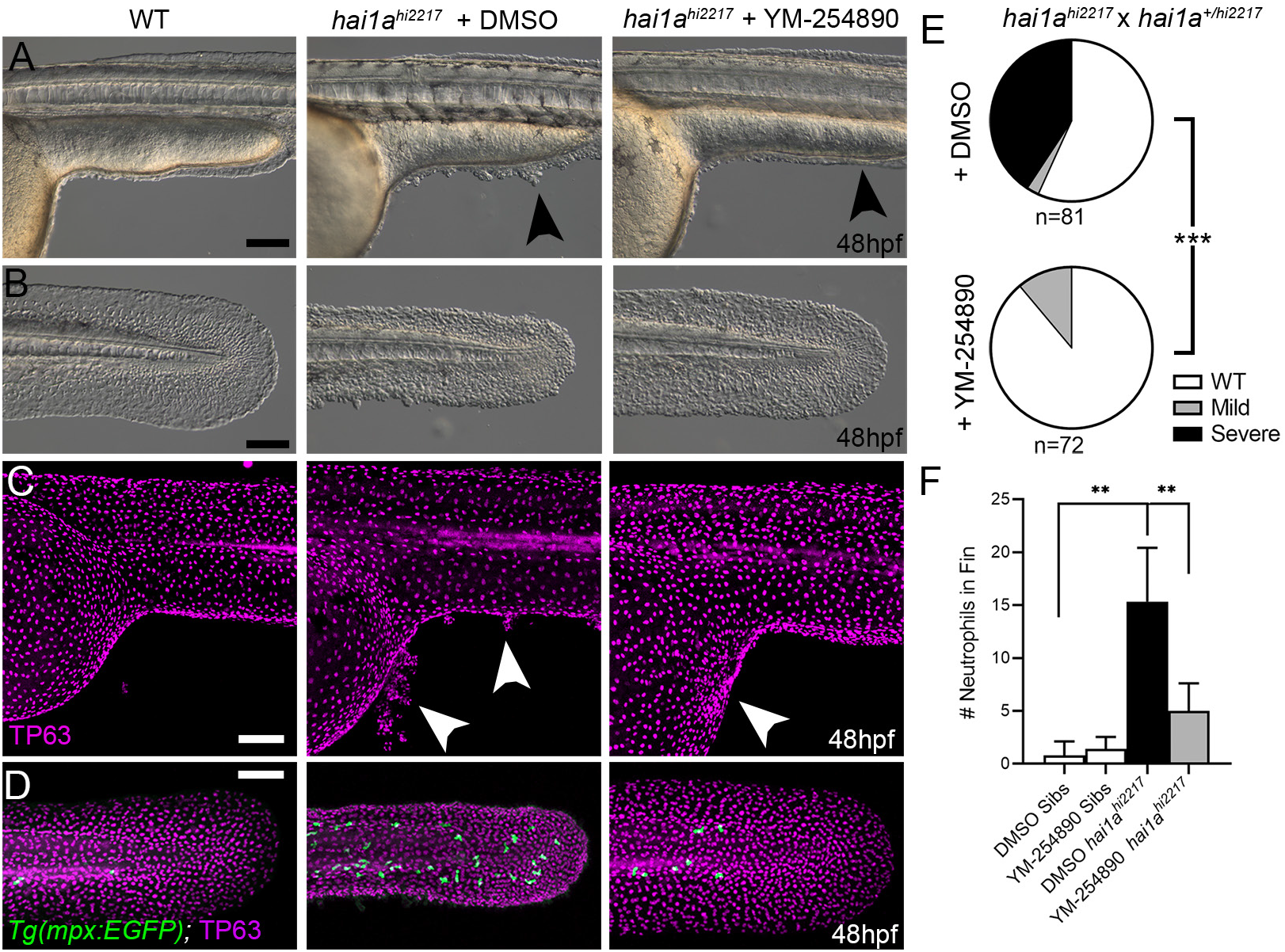
Gq inhibition rescues both epidermal and inflammation phenotypes of *hai1a* mutants. **A-D:** Lateral images of ventral trunk and tail at 48hpf for WT (left panels), *hai1a*^*hi2217*^ treated with 0.5%DMSO (middle panels) and *hai1a-*^*hi2217*^ treated with 32μM YM-254890 (right panels). DIC micrographs are shown in A-B, whilst projected confocal images are shown in C-D, where embryos are immunostained for TP63 (C, D; magenta) and eGFP (D; green). Embryos in (D) are hemizygous for *Tg(mpx:eGFP)*^*i114*^. Arrowheads indicate region of aggregate formation lost upon treatment with Gq inhibitor YM-254890. **E:** Pie charts showing proportion of embryos with no (WT; white), mild (grey) or severe (black) *hai1a* mutant epidermal phenotypes. Embryos were derived from *hai1ahi2217/hi2217* x *hai1a+/hi2217* crosses and assayed at 48hpf. Clutches treated with 0.5% DMSO (upper pie chart) were compared to those treated with 32μM YM-254890 (lower pie chart) by Chi-squared analysis. *** = p<0.001; n=72. **F:** Graph of counts of eGFP positive neutrophils in the fins at 48hpf of *Tg(mpx:EGFP)*^*i114*^, or *hai1a*^*hi2217*^; *Tg(mpx:EGFP)*^*i114*^ treated with 0.5% DMSO, or 32μM YM-254890. n= 6; Mann-Whitney test; ** = p<0.01. Scale bars A, B, C, D = 100μm.

### PMA treatment phenocopies the *hai1a* mutant

As IP_3_R inhibition only blocks inflammation in *hai1a* mutants, but an inhibitor of a PLC activator (Gq) additionally reduces the epidermal defects, we considered that diacyl glycerol (DAG) might contribute to the epidermal defects as the second product of PIP_2_ cleavage (along with IP_3_). Indeed, treating zebrafish embryos with phorbol 12-myristate 13-acetate (PMA), a DAG analogue, has been described to generate global embryological phenotypes (Hrubik et al., 2016). Treating WT embryos from 15hpf to 24hpf with 125ng/ml PMA resulted in embryos with striking similarities to strong *hai1a* mutants, including a thin or absent yolk sac extension, lack of head straightening, lack of lifting the head off the yolk, and multiple epidermal aggregates on the skin (Fig. 6A-C). These aggregates were due, at least partially, to displacement of basal keratinocytes as shown by TP63 staining where the basal keratinocyte nuclei lost their uniform distribution (Fig. 6D-E). Treatment of WT embryos from 24hpf to 48hpf with 125ng/ml PMA led to a fin defect similar to the dysmorphic *hai1a* mutant fin (Fig. 6F-G). It has been shown that the basal keratinocytes in *hai1a* lose their epithelial nature and adopt a partially migratory phenotype (Carney et al., 2007) (Video 5). We treated *Tg(krtt1c19e:lyn-tdtomato)*^*sq16*^ larvae (Lee, Asharani, & Carney, 2014) with 37.5ng/ml PMA for 12 hours and imaged the basal epidermis at 3dpf by light-sheet timelapse. Whilst the DMSO treated transgenic larvae had very stable keratinocyte membranes and shape, PMA treatment led to a less stable cell membrane topology and dynamic cell shape, similar to *hai1a* mutants (Fig. 6H; Videos 5, 6). Kymographs taken from Video 6 highlighted both the more labile and weaker cell membrane staining following PMA treatment (Fig. 6I). The potency of PMA was dependant on region and reduced with age

**Figure 6:**
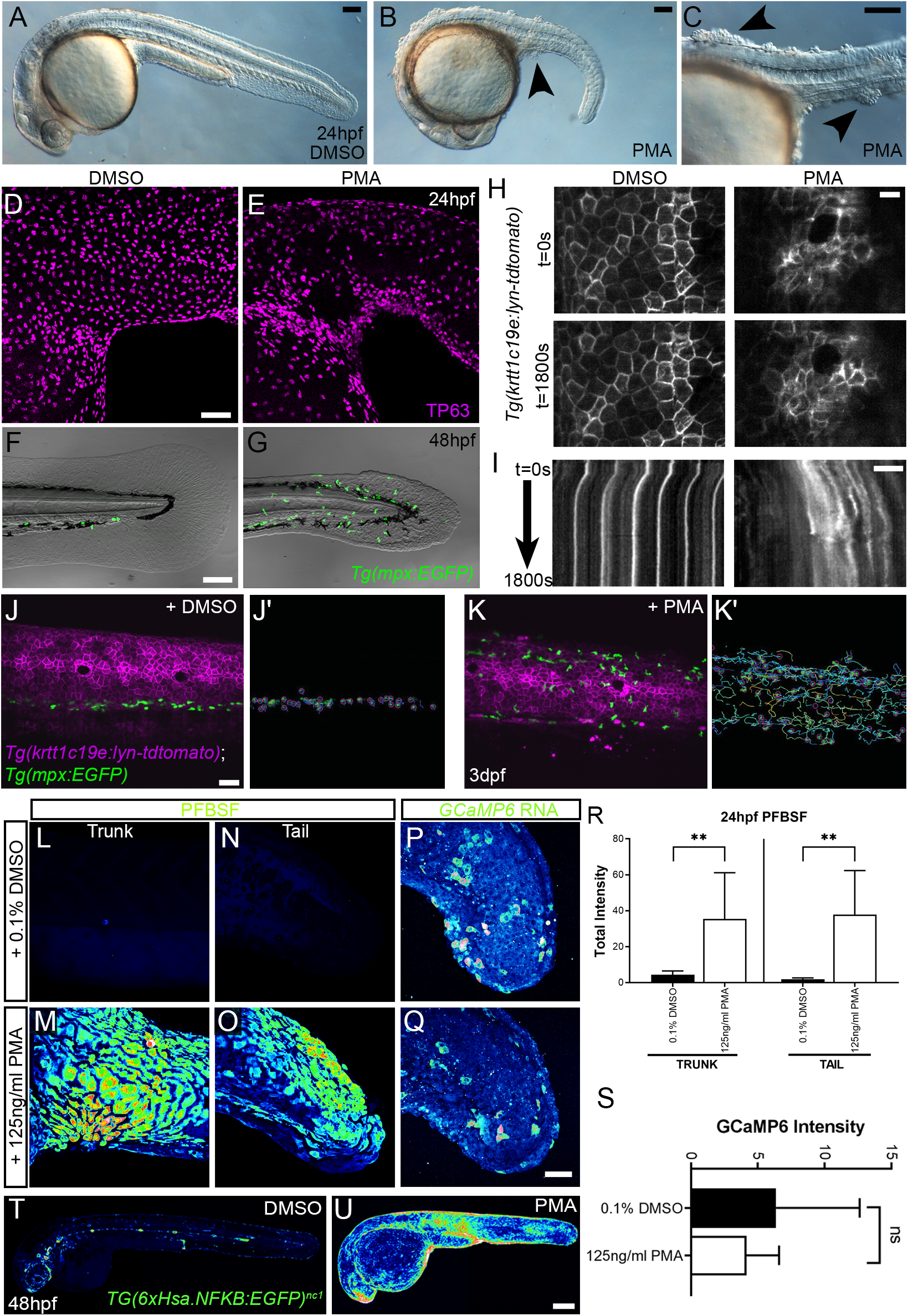
PMA induces epidermal aggregates, motility, H_2_O_2_, NfkB and inflammation. **A-B:** Lateral micrographs of embryos treated with DMSO (A) or 125ng/ml PMA (B, C) showing generation of epidermal aggregates (arrowheads). **D-E:** Projected confocal images of the trunk of 24hpf WT embryos treated with 0.1% DMSO (D) or 125ng/ml PMA (E) and immunostained for TP63 (magenta), showing aggregation of TP63 positive cells. **F-G:** Projected confocal images superimposed on DIC image of the tail of 48hpf *Tg(mpx:EGFP)*^*i114*^ embryos treated with 0.1% DMSO (F) or 125ng/ml PMA (G) showing fin defect and activation of eGFP positive neutrophils (green, G). **H-I:** Single timepoint images at t=0 (top panels, H) and t=1800s (lower panels, H) and kymographs (I) derived from light-sheet movies (Video 6) of the epidermis of 3dpf *Tg(krtt1c19e:lyn-tdtomato)*^*sq16*^ larvae treated with 0.1% DMSO (left panels) or 37.5ng/ml PMA (right panels) showing the lack of membrane stability following PMA treatment. **J-K’:** Single frames (J, K) and tracks of eGFP positive neutrophils (J’, K’) from light-sheet Video 7 showing neutrophils labelled by eGFP and basal keratinocyte cell membranes labelled by lyn-tdTomato in the trunk of a 3dpf *Tg(krtt1c19e:lyn-td-tomato)*^*sq16*^ larva treated with 0.1% DMSO (J, J’) or 37.5ng/ ml PMA (K, K’) for 18 hours, and imaged every 20 sec for 30 minutes Track colour in J’, K’ denotes mean velocity (dark blue 0.0 – red 0.2). **L-O:** Projected lateral confocal views of PFBSF staining of 24hpf WT embryos treated with 0.1% DMSO (L, N) or 125ng/ml PMA (M, O) showing elevation of H_2_O_2_ in the trunk (L, M) and tail (N, O). **P-Q:** Projected confocal images of eGFP in the tail at 24hpf of WT injected with *GCaMP6s* RNA, treated with DMSO (P), or with 125ng/ml PMA (Q). Images are temporal projections of timelapse movies taken at maximum speed intervals (2 minutes) and projected by time. **R:** Plot of PFBSF fluorescent staining intensity of WT embryos treated with 0.1% DMSO or 125ng/ml PMA in the trunk and tail. n=6; ANOVA with Bonferroni post-test ** p<0.01. **S:** Graph comparing eGFP intensities from 24hpf *GCaMP6s* RNA timelapses in tail following treatment with DMSO and 125ng/ml PMA. n= 10; t-test. **T-U:** Lateral confocal projections of *Tg(6xHsa.NFKB:EGFP)nc1* embryos reporting NfkB signalling levels at 48hpf in WT treated with DMSO (T), and WT treated with 125ng/ml PMA (U). Scale bars A, B, C, F= 100μm; D, J, Q = 50μm; H, I = 20μm; U = 200μm.

Most PMA treated *Tg(mpx.eGFP)*^*i114*^ larvae at 48hpf also had more neutrophils in the epidermis than untreated controls, which were highly migratory (Fig. 6F-G, J-K’; Video 7). We determined H_2_O_2_ levels in PMA treated embryos using PFBSF staining, and found it was significantly increased in both trunk and tail at 24hpf (Fig. 6 L-O, R). In contrast, when we treated *GCaMP6s* RNA injected embryos with PMA, we failed to see an increase in calcium flashes, as seen in *hai1a* (Fig. 6P, Q, S). To see if the heightened H_2_O_2_ and inflammation was also correlated with increased NfkB signalling we treated *Tg(6xHsa.NFKB:EGFP)*^*nc1*^ embryos with 125ng/ml PMA. There was a robust increase in fluorescence indicating that PMA activates the NfkB pathway (Fig. 6T, U).

The phenocopy and the rescue of *hai1a* by PMA and Gq inhibition respectively implies DAG is elevated in *hai1a* mutants. If this is true, then inhibition of its target, Protein Kinase C, should also ameliorate the *hai1a* mutant phenotype. We treated *hai1a*^*hi2217*^ embryos with the PKC inhibitor, GFX109203, and found this was able to reduce the epidermal aggregates and disruption of fin morphology as imaged by DIC or immunostaining for TP63 (Fig. 7A-F). Neutrophil inflammation in the epidermis was somewhat reduced, but not to a significant degree (Fig. 7C-G). Thus, the epithelial defects of *hai1a* are due to PKC activation.

**Figure 7:**
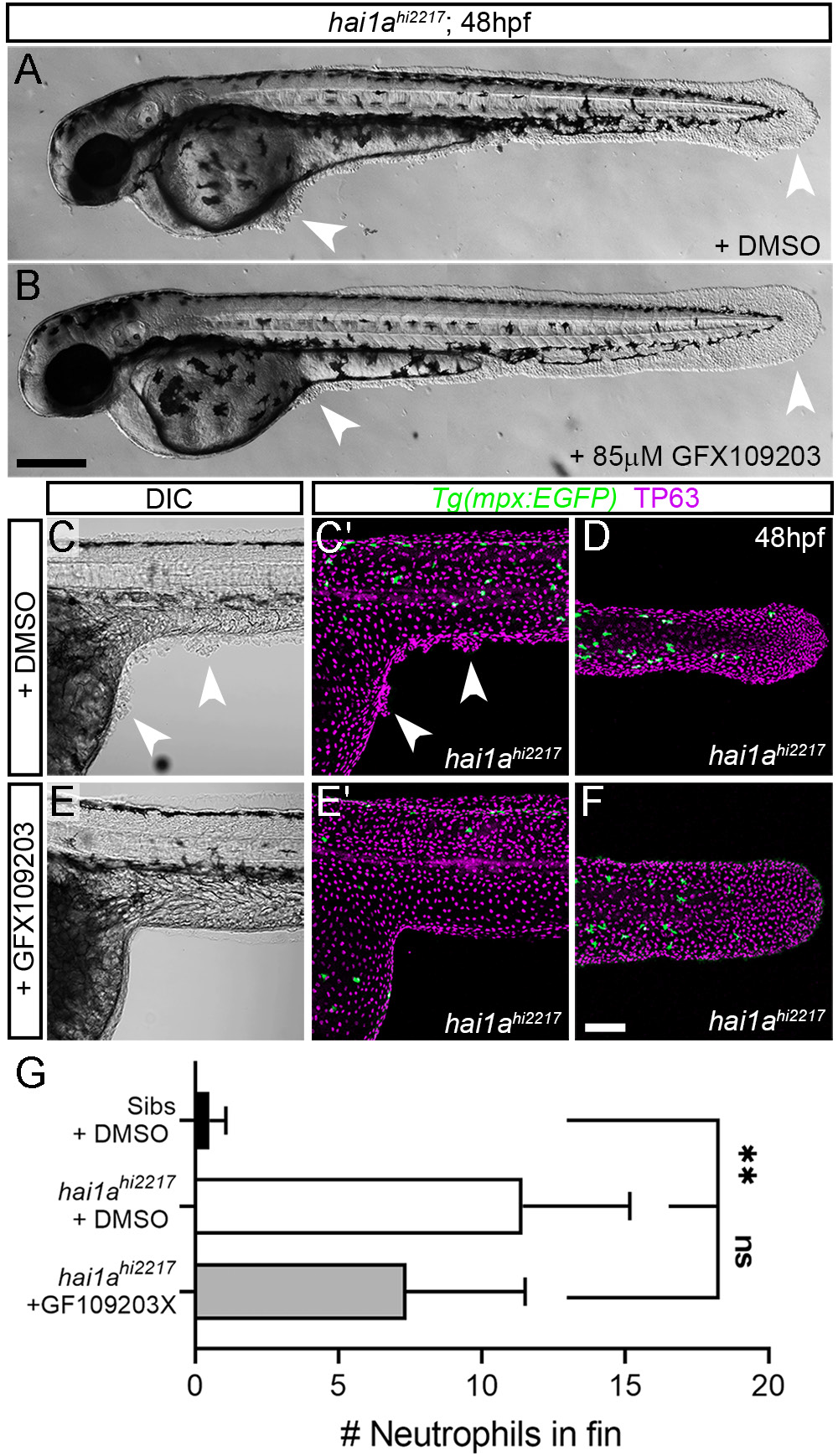
Inhibition of PKC rescues epidermal defects of *hai1a*. **A-B:** Lateral brightfield images of 48hpf *hai1a*^*hi2217*^ larvae treated with 0.5% DMSO (A) or 85μM GFX109203 (B). Epidermal aggregates and fin deterioration are rescued by the PKC inhibitor (arrowheads). **C-F:** DIC (C, E) and projected confocal images (C’, E’, D, F) of *hai1a*^*hi2217*^; *Tg(mpx:EGFP)*^*i114*^ trunk at 24hpf(C-C’, E-E’) and tail at 48hpf (D, F), either treated with 0.5% DMSO (C-D) or 85μM GFX109203 (E-F). Embryos are immunostained for TP63 (magenta) and eGFP (green), highlighting rescue of epidermal phenotype and partial rescue of neutrophils by GFX109203. **G:** Counts of eGFP positive neutrophils in the fins at 48hpf of *Tg(mpx:EGFP)*^*i114*^, or *hai1a*^*hi2217*^; *Tg(mpx:EGFP)*^*i114*^ treated with 0.5% DMSO, or 85μM GFX109203. n= 8; ANOVA, Dunn’s Multiple comparisons; ** = p<0.01. Scale bars B = 200μm; F = 100μm.

### Elevated MAPK signalling generates epithelial defects in hai1a

We next sought to determine which pathways downstream of PKC are responsible for the epidermal defects. The MAPK pathway is a major target pathway of multiple PKC isoforms, and activation of this pathway in zebrafish epidermis has previously been shown to induce papilloma formation which have very similar attributes to *hai1a* mutant aggregates (Chou et al., 2015). Although whole embryo western analysis of *hai1a* mutants failed to show an overall increase in pERK (Armistead et al., 2020), we performed wholemount immunofluorescent analysis in case there was only a localized effect. Indeed, we observed a significant and localised increase in cytoplasmic pERK immunoreactivity (phospho-p44/42 MAPK (Erk1/2) (Thr^202^/Tyr^204^)) in the regions of epidermal aggregate formation in *hai1a* mutants and in PMA treated embryos, including under the yolk at 24hpf and in the fins at 24hpf and 48hpf (Fig. 8A-H, M; Fig. S5A-F). There was no increase in total ERK levels in the mutant (Fig. 8K-L). Increased pERK was seen in the both the cytoplasm and nucleus of TP63 positive cells but was only increased in the nucleus of periderm cells (Fig. 8F; Fig. S5D). To establish that this is an early marker of aggregate formation, and not a sequala, we stained *hai1a* mutant embryos at earlier timepoints. We found that even nascent aggregates at 16hpf have pERK staining (Fig. 8I-J), which increases in number over time (Fig. S5G-J).

**Figure 8:**
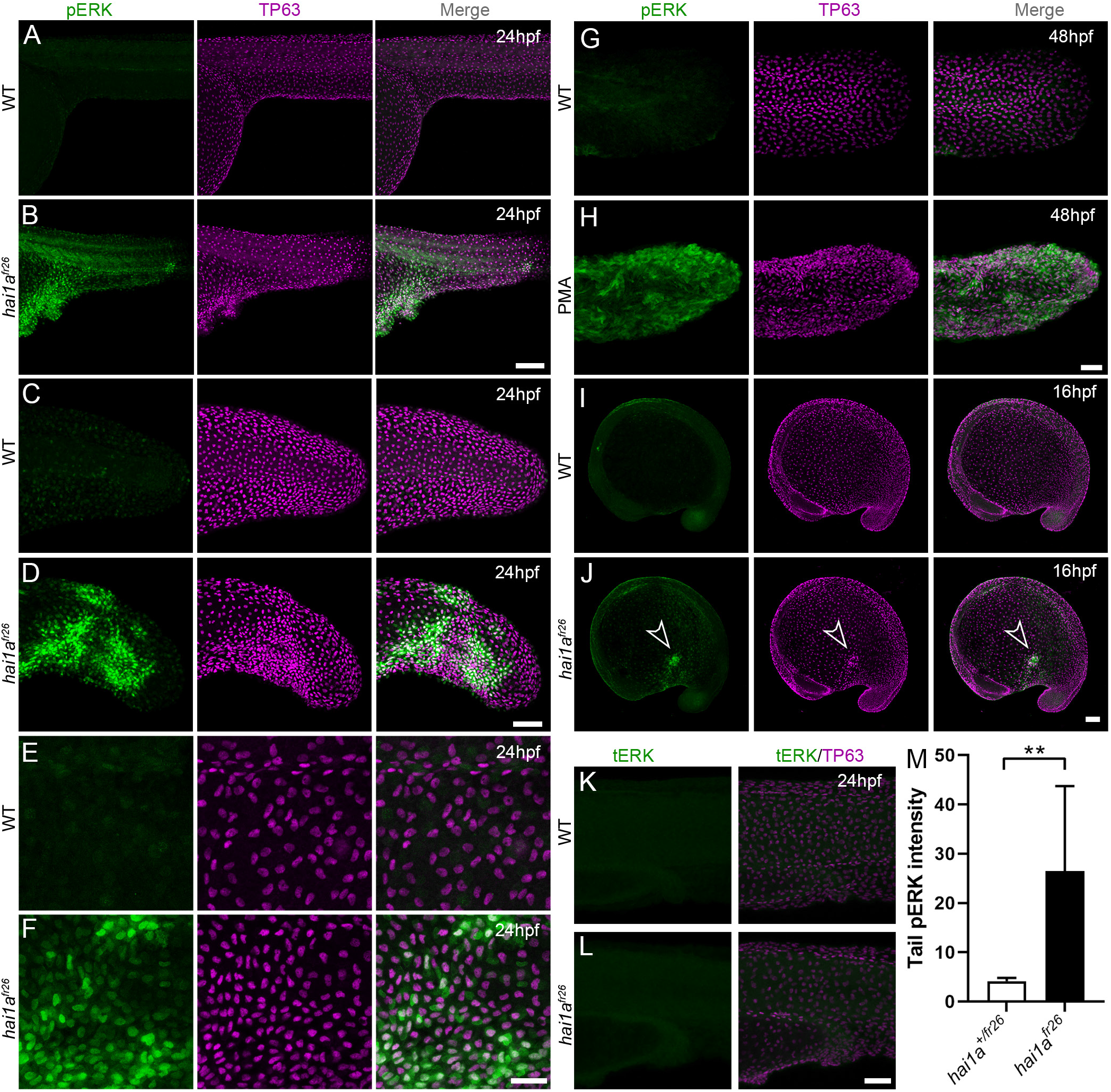
Elevation of pERK levels in PMA treated and *hai1a* mutant epidermis. **A-L:** Lateral projected confocal images of trunks (A, B, E, F, K, L), and tails (C, D, G, H) of embryos immunostained for TP63 (A-L; magenta) and pERK (green; A-J) or total ERK (tERK; green K, L) at 24hpf (A-F, K, L), 48hpf (G, H) and 16hpf (I, J). Both *hai1a*^*fr26*^ (B, D, F, J) and 125ng/ml PMA treated (H) embryos show increased epidermal pERK levels compared to untreated WT (A, C, E, G, I). There was no increase in tERK in *hai1a*^*fr26*^ (L) compared to WT (K). Elevation of epidermal pERK in *hai1a*^*fr26*^ mutants and PMA treated embryos is seen in the trunk (B, F) and tail (D, H) as well as in nascent aggregates (arrowhead, J). **M:** Quantification of pERK immunofluorescent intensity in the tail of 24hpf *hai1a*^*fr26*^ larvae compared to siblings. n= 5; Mann-Whitney test; ** = p<0.01. Scale bars B, J = 100μm; D, H, L = 50μm; F = 20μm.

To determine if elevated pERK is causative of epidermal defects, we attempted to rescue using pERK inhibitors. Initially we used PD0325901, however this appeared to give fin fold deformities, even in WT embryos (Anastasaki, Rauen, & Patton, 2012), precluding ability to assess rescue in *hai1a*, although there was a noticeable reduction in epidermal aggregates forming under the yolk-sac extension (data not shown). Instead we tried U0126 and CI-1040, other well-known pERK inhibitors (Allen, Sebolt-Leopold, & Meyer, 2003; Favata et al., 1998). Both inhibitors showed a significant reduction in *hai1a* mutant epidermal aggregates under the yolk, and restoration of the overall and tail epithelial morphology, with embryos showing a *hai1a* phenotype class significantly reduced (Fig. 9A-G; Fig. S6A-F). Similarly, the epidermal defects of the trunk, yolk and tail following PMA treatment were also ameliorated by concomitant U0126 treatment (Fig. 9H-I; Fig. S6G-H). Rescue of aggregates and tail morphology following PMA treatment or in *hai1a* mutants could be visualized by immunolabelling TP63 in basal keratinocyte nuclei (Fig. 9J-O; Fig. S6I-J). Treatment with U0126 did not significantly reduce neutrophil inflammation of *hai1a* mutants or PMA treatment (Fig. 9L-P). It has been shown that the epidermal defects in *hai1a* are associated with loss of E-Cadherin from adherens junctions (Carney et al., 2007). As there was a rescue of the epithelial phenotype following pERK inhibition, we looked at the status of the adherens junction marker βcatenin. Whilst the WT basal epidermal cells of the 48hpf tail showed strong staining at the membrane, *hai1a* mutants and PMA treated embryos showed a significant loss of βcatenin at the membrane and increase in the cytoplasm (Fig. 9Q-V, Y, Z). Treatment of *hai1a* mutants with U0126 restored the membrane localisation of βcatenin (Fig. 9W, X, AA).

**Figure 9:**
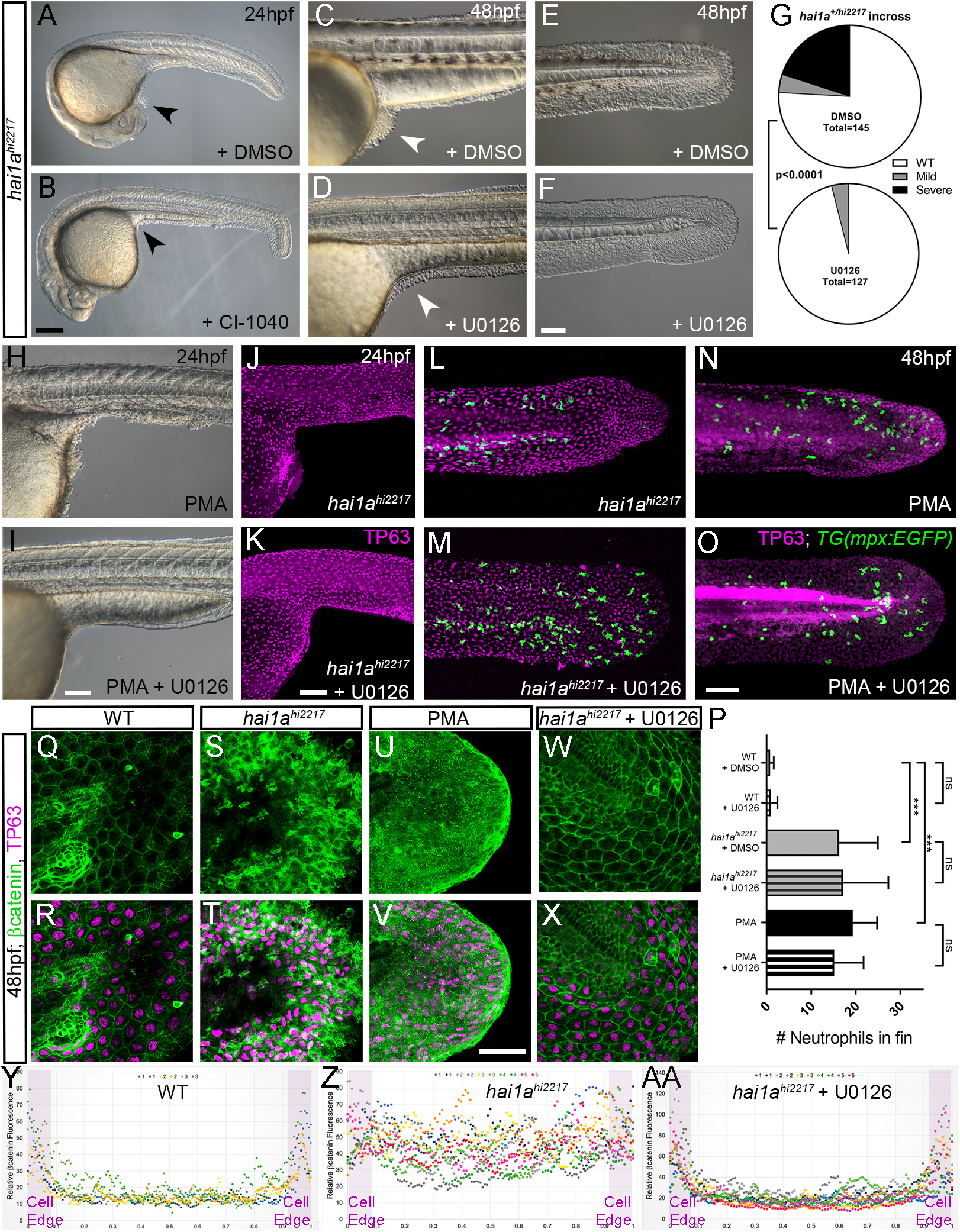
Rescue of the *hai1a* epidermal phenotype by pERK inhibitors. **A-F:** Lateral DIC images of 24hpf (A-B) or 48hpf (C-F) *hai1a*^*hi2217*^ embryos treated with either DMSO (A, C, E), CI-1040 (B) or U0126 (D, F) showing rescue of general morphology (B), trunk (D) and tail (F) epidermal phenotypes compared to DMSO treated *hai1a*^*hi2217*^. Epidermal aggregates under the yolk are reduced in the treated mutants (A-D; arrowheads). **G:** Proportion of 48hpf larvae derived from *hai1a*^*+/hi2217*^ in-cross showing mild or severe *hai1a* epidermal phenotype following DMSO (upper) or U0126 (lower) treatment (Chi-squared test). **H-I:** Lateral DIC images of 24hpf embryo treated with 125ng/ml PMA (H) or PMA and U0126 (I). Yolk associated epidermal aggregates are reduced. **J-M:** Lateral projected confocal images of *hai1a*^*hi2217*^; *Tg(mpx-:eGFP)*^*i114*^ trunk at 24hpf(J-K) and tail at 48hpf (L-M), either treated with 0.5% DMSO (J, L) or U0126 (K, M). Embryos are immunostained for TP63 (magenta) and eGFP (green), highlighting rescue of epidermal phenotype but no reduction of neutrophils. **N-O:** Lateral projected confocal images of *Tg(mpx-:eGFP)*^*i114*^ treated with PMA alone (N) or PMA with U0126 (O) and immunostained for TP63 (magenta) and eGFP (green). Fin morphology is restored but neutrophils are still present. **P:** Quantification of neutrophils in the fins showing U0126 does not reduce inflammation induced by loss of *hai1a* or PMA treatment. n=8; ANOVA with Bonferroni post-test; ***=p<0.001. **Q-X:** Projected confocal images of 48hpf larval tails immunostained for βcatenin (green) and TP63 (magenta; R, T, V, X) of WT (Q, R, U, V) and *hai1a*^*hi2217*^ (S, T, W, X), either untreated (Q-T), treated with PMA (U, V) or U0126 (W, X). **Y-AA:** Profile plots of fluorescence distribution across cells of WT (Y), *hai1a*^*hi2217*^ (Z) and *hai1a*^*hi2217*^ treated with U0126 (AA). X-axis represents width of the cell. βcatenin immunofluorescence intensity (Y-axis) shows majority at cell edge (demarcated in light purple) in WT and rescued *hai11a* mutants, but is distributed in cytoplasm in mutant. 2 cells per 3-5 larvae were analysed. Scale bars: B = 200μm, F, I, K, O = 100μm, V = 20μm.

### Phosphorylation of cytoplasmic RSK by pERK, leads to loss of E-cadherin at the *hai1a* keratinocyte membrane

As increased pERK appeared to contribute strongly to loss of adherens junctions, and removal of E-Cad-herin/βcatenin from the membrane, we sought to determine how pERK signalling might affect adherens junctions. We predicted that this would occur through a cytoplasmic target of pERK, as we have previously shown that there is no transcriptional downregulation of E-cadherin levels in *hai1a,* making a nuclear transcription factor target less likely to be relevant. The p90RSK family of kinases represent cytoplasmic direct targets of Erk1/2 phosphorylation which regulate cell motility, and thus were good candidates for mediators disrupting cell-cell adhesion (Caslavsky, Klimova, & Vomastek, 2013; Tanimura & Takeda, 2017). We determined that at least RSK2a (=p90RSK2a, encoded by *rps6ka3a*) is expressed in the basal keratinocytes at 24hpf (Fig. 10A, B), and other RSK family members may also be expressed there. To gauge if there was an alteration in phosphorylation of RSK family members in the epidermis of *hai1a* mutants, we used an anti-body which detects a phosphorylated site of mouse p90RSK (Phospho-Thr^348^). This site is phosphorylated in an ERK1/2 dependant manner (Romeo, Zhang, & Roux, 2012). We noticed a substantial increase in cytoplasmic signal in both *hai1a* mutants and PMA treated embryos. Where p90RSK-pT^348^ signal was largely nuclear in both basal and periderm cells in WT, it was more broadly observed in *hai1a* mutant fins, with an increase in the cytoplasm leading to a more uniform staining (Fig. 10C-D’). This cytoplasmic signal was lost upon U0126 treatment showing it was pERK dependant (Fig. 10E-E’). Similarly, increased cytoplasmic p90RSK-pT^348^ was observedfollowing PMA treatment that was reduced by co-treatment with U0126 (Fig. 10F-H’). The increase in cytoplasmic p90RSK-pT^348^ signal, and its reduction by U0126, was significant in both *hai1a* mutants and PMA treated embryos (Fig. 10I-J).

**Figure 10:**
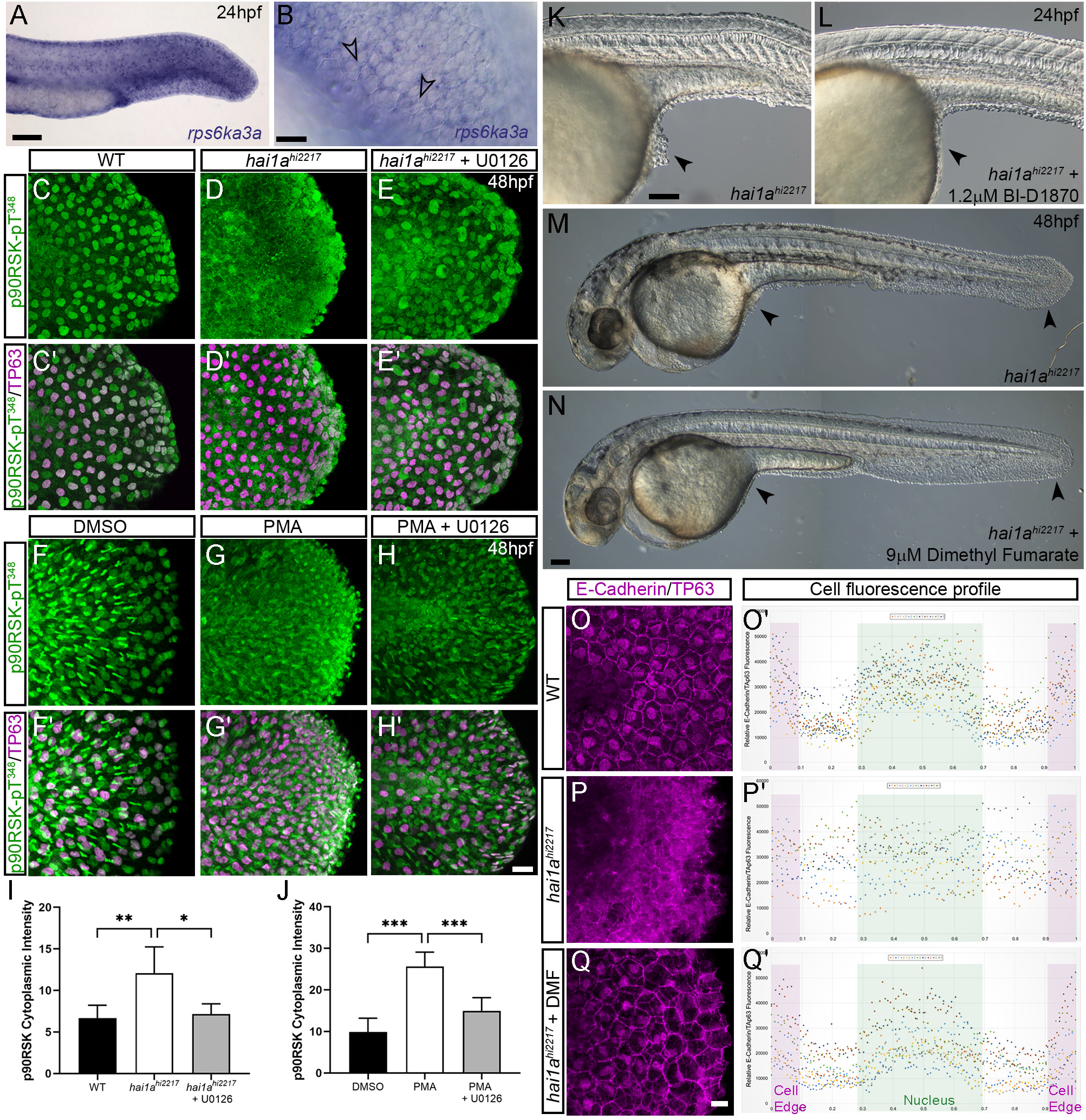
Altered RSK status in *hai1a*^*hi2217*^ accounts for epidermal defects. **A-B:** In situ hybridisation of *rps6ka3a* at 24hpf under low (A) and high (B) power magnification showing expression in basal keratinocytes. Open arrowheads in B indicate borders of EVL cells bisecting nuclei of underlying *rps6ka3*a positive cells. **C-H’:** Lateral projected confocal images of the tails of embryos immunostained for p90RSK (Phospho-Thr^348^) (C-H’) and TP63 (C’-H’). In both the *hai1a*^*hi2217*^ (D-D’) and 125ng/ml PMA treated (G-G’) embryos, there is an increase in cytoplasmic levels of p90RSK (Phospho-Thr^348^) signal above the nuclear only signal seen in WT (C-C’) or DMSO (F-F’). Treatment with the pERK inhibitor U0126 reduced cytoplasmic levels but did not affect nuclear signal (E-E’, H-H’), **I-J:** Quantification of immunofluorescent intensity of cytoplasmic levels of p90RSK (Phospho-Thr^348^) in basal keratinocytes of tails of 48hpf WT and *hai1a*^*hi2217*^, treated with DMSO or U0126 (I), and PMA or PMA plus U0126 (J). Nucleus signal was excluded by masking from the DAPI channel. n=5; t-test; *** = p<0.001, ** = p<0.01, * = p<0.05. **K-N:** Lateral DIC images of *hai1a*^*hi2217*^ embryos at 24hpf (K, L) and 48hpf (M, N) untreated (K, M) or treated with 1.2μM BI-D1870 (L) or 9μM Dimethyl Fumarate. Locations of epidermal aggregates and loss of tail fin morphology in *hai1a* mutants, and their rescue by RSK inhibitor treatment are indicated by arrowheads. **O-Q:** Lateral projected confocal images of the tails of embryos immunostained with antibodies against E-Cadherin and TP63in WT (O), (P) and *hai1a*^*hi2217*^ treated with DMF (Q). **O’-Q’:** Profile plots of fluorescence distribution across cells of WT (O), *hai1a*^*hi2217*^ (P’) and *hai1a*^*hi2217*^ treated with DMF (Q’). X-axis represents width of the cell. βcatenin immunofluorescence intensity (Y-axis) shows majority at cell edge (E-Cadherin domain demarcated in light purple) and centre of cell (Nucleus demarcated in light green) in WT and rescued *hai11a* mutants, but there is no clear membrane signal in the untreated *hai1a* mutants. 2 cells per 5 larvae were analysed. Scale Bars: A, K, N = 100μM; B, H, P = 20μM.

If phosphorylation of an RSK protein is required for mediating the pERK epidermal defects in *hai1a* mutants, then inhibition of RSK should rescue the epidermal defects. As morpholino targeted inhibition of *rps6ka3a* was unsuccessful, we employed established pan-RSK inhibitors BI-D1870 and Dimethyl Fumarate (Andersen et al., 2018; Sapkota et al., 2007). We noted that both inhibitors were able to reduce epidermal aggregates in *hai1a* mutants and restore fin morphology, when visualised by DIC or TP63 immunofluorescence (Fig. 10K-N; Fig.S7 A-B, D-E). Reduction of mutant phenotype classes was significant at both 24hpf and 48hpf (Fig. S7C). We then assayed if RSK inhibition can reduce the aberrant cytoplasmic E-Cadherin staining in *hai1a* mutant basal keratinocytes and observed that Dimethyl fumarate treatment restored membrane localisation of E-Cadherin in the mutants (Fig. 10O-Q’). Thus, phosphorylation of RSK proteins is altered in *hai1a* mutants and their inhibition can restore E-Cadherin to the membrane and reduce epidermal aggregate formation.

## Discussion

There are a number of similarities between loss of Hai1a in zebrafish and overexpression of Matriptase in the mouse epidermis, including inflammation, hyperproliferation and enhanced keratinocyte motility. This implies that there is an ancient function for this protease/protease inhibitor system. In mouse and human, the loss of Matriptase causes stratum corneum defects including ichthyosis, loss of filaggrin processing and defects in desquamation. As there is no stratum corneum in zebrafish, this cannot be the ancestral role. Since the only overt phenotype in zebrafish Matriptase1a maternal zygotic mutants was transient loss of ear otoliths, and *hai1a*; *st14a* double mutants appear normal, it is unclear why the Hai1-Matrip-tase system evolved in fish at all. Our previous analysis of the epithelial aggregates in zebrafish *hai1a* mutants led us to conclude that nascent aggregates are not forming due to increased proliferation, which occurs at later timepoints, but due to increase keratinocyte motility. Further, although early and prominent, we have shown that the inflammation in *hai1a* mutants is not required for epithelial defects but occurs in parallel to it. These two primary phenotypes, sterile inflammation and epithelial cell motility, are the hallmarks of the damage response during early wound detection. Wounds are detected through a combination of osmotic surveillance, release of DAMPs from cell lysis and mechanotransduction at the wound edge (Enyedi & Niethammer, 2015). Indeed, disrupted osmolarity differences and nucleotide release in the zebrafish epidermis generate inflammation and epithelial cell motility as part of a damage response (de Oliveira et al., 2014; Enyedi & Niethammer, 2015; Gault, Enyedi, & Niethammer, 2014; Hatzold et al., 2016), with the resulting phenotypes strikingly similar to *hai1a* mutants. Further, PAR2 promotes cell migration in scratch assays in conjunction with P2Y purinergic and EGF receptors (K. Shi, Queiroz, Stap, Richel, & Spek, 2013). It is, therefore, tempting to propose the Hai1-Matriptase system as part of the early wound detection process. How it might act as a sensor is not clear; however, Matriptase activity is highly sensitive to reduced pH and has been linked to response to acidosis (Tseng et al., 2010). Altered pH has recently been proposed to regulate wound progression (Stolwijk et al., 2020). In the mammalian epidermis, Matriptase becomes active at the transition to the stratum corneum where the matrix becomes acidic (Miller & List, 2013), and the acid sensing ion channel, ASIC1, is also a target of Matriptase further supporting a link between Matriptase and environmental pH (Clark, Jovov, Rooj, Fuller, & Benos, 2010). Matriptase activation is also increased by redox state and the oxidative effects of heavy metals such as cadmium and cobalt, and loss of Matriptase makes cells more sensitive to heavy metal toxicity (J. K. Wang et al., 2014). We postulate that the Hai1-Matriptase system evolved in the epidermis as a surveillance system of altered external pH, osmolarity, redox state or metal ions, which may arise due to wounding of the epithelial barrier. Such a surveillance system would likely act redundantly with other pathways of the damage response, and promote responses through PAR2 signalling as also suggested by Schepis et al. (2018).

There are strong links between Matriptase dysregulation and cancer progression (Martin & List, 2019), and the cellular and tissue level phenotypes of *hai1a* have similarities to tumours. Tumours have long been considered to represent non-healing wounds and transformed cells in zebrafish both promote and respond to inflammation through similar mechanisms to wound responses (Feng, Santoriello, Mione, Hurlstone, & Martin, 2010; Schafer & Werner, 2008). We have shown that the inflammation in *hai1a* has similar molecular signatures to that following wounding, or other damage sensors such as ATP release and osmotic stress. Indeed, we show that increased IP_3_R-dependant calcium release in *hai1a* epidermis activates Duox, leading to high hydrogen peroxide levels, which in turn leads to increased NfkB signalling. This pathway has also been shown to act downstream of ATP via purinergic GPCRs, to recruit neutrophils in zebrafish (de Oliveira et al., 2014). We were unable to definitively rescue *hai1a* phenotypes with a PLC inhibitor due to toxicity, however an inhibitor of Gq very effectively rescued the inflammation, but surprisingly also the epithelial defects. We showed that the DAG analogue, PMA can phenocopy the epidermal defects of *hai1a* mutants. Conversely, inhibiting the DAG receptor, PKC, rescued the epithelial phenotypes. Furthermore, PMA treatment increased H_2_O_2_, NfkB and neutrophil inflammation, which is in line with known activation of Duox and IKK by PKC (Rigutto et al., 2009; Turvey et al., 2014). Seminal experiments in transgenic mice overexpressing Matriptase in the epidermis and treated with a DMBA/ PMA regime, concluded that Matriptase and PMA activate functionally similar carcinoma promoting pathways (List et al., 2005). Our subsequent analysis suggests that this would include the MAPK pathway, as we see increased phosphorylated-ERK in the epidermis of both *hai1a* mutants and also PMA treated embryos. This was surprising as previous reports did not detect this by western analysis of whole embryos, although we observed pERK increased only in the epidermis, which may account for a limited global proportion. That we can rescue the epithelial defects using a MEK inhibitor indicated that this increase in epidermal pERK is critical to the phenotype. The MAPK pathway, in addition to well-defined roles in cell proliferation, translation, differentiation and survival, is known to regulate cell motility (Tanimura & Takeda, 2017). Indeed, misexpression of activated MEK2 in zebrafish epidermis generated papillomas with remarkable resemblance to the epidermal aggregates in *hai1a* mutants (Chou et al., 2015), and which are not overtly proliferative. Additionally, PAR2 activates motility of an oesophageal tumour line partially via MAPK/ERK pathway (Sheng et al., 2019). One major effector phosphorylated by pERK is the kinase RSK2 (= p90RSK2), which is encoded by the *RPS6KA3* gene. Like Matriptase, activation of RSK2 is associated with tumour progression (Kang & Chen, 2011), particularly promoting invasiveness and metastasis of glioblastomas and head and neck squamous cell carcinomas (Kang et al., 2010; Sulzmaier et al., 2016). Promotion of invasiveness has also been noted for other members of the p90RSK family such as RSK1, activation of which promotes invasion of melanoma clinically as well as *in vitro* and zebrafish melanoma models (Salhi et al., 2015). MAPK signalling has also been shown to reduce E-Cadherin expression at adherens junctions and promoting its cytoplasmic accumulation in an RSK dependant manner, without reducing E-Cadherin protein levels (Caslavsky et al., 2013). We previously showed an identical effect on E-Cadherin relocation in zebrafish *hai1a* mutants (Carney et al., 2007). Intriguingly, recent proximity protein labelling has identified p120-catenin being phosphorylated by p90RSK activity. This catenin promotes cell-cell adhesion by regulating Cadherin stability at the membrane, a function inhibited by p90RSK phosphorylation (Meant et al., 2020). More broadly, RSK2 activity promotes cell motility through other mechanisms, including inactivation of Integrins and activation of the RhoGEF, LARG (Gawecka et al., 2012; G. X. Shi, Yang, Jin, Matter, & Ramos, 2018). Thus, we propose pERK signalling, through p90RSK members, significantly contributes to the *hai1a* epidermal phenotype.

Our analyses allow us to propose a pathway which accounts for both the inflammatory and the epidermal phenotypes (Fig. 11). Whilst the inflammatory phenotype is strongly reliant on IP_3_ and calcium release, there is likely a partial contribution from PKC activation, as PMA also promotes inflammation and PKC inhibition partially reduces inflammation. In contrast, only inhibition of Gaq, PKC, pERK and p90RSK rescued the epidermal phenotype. This model invokes activation of Gq and Phospholipase C to generate Ca^++^ and DAG. This has been well documented to occur downstream of PAR2 in many cell types including keratinocytes, where inhibition of Gq and PKC reduces PAR2 mediated Nfκb signalling (Bohm et al., 1996; Goon Goh et al., 2008; Macfarlane et al., 2005). PAR2 agonists have also been shown to activate the ERK1/2 MAPK pathway, although the mechanisms proposed are varied. In breast cancer cell lines, ERK is activated by PAR2 and promotes invasiveness and correlates with PAR2 mediated PIP2 hydrolysis (Jiang et al., 2004; Morris et al., 2006). Similarly, PAR2 stimulation of astrocyte migration required Gq, but not Gi/o, stimulation of pERK (McCoy, Traynelis, & Hepler, 2010). However, a number of publications have implicated EGFR transactivation in signalling downstream of PAR2, including stimulation of the MAPK pathway (Darmoul, Gratio, Devaud, & Laburthe, 2004). Other work has indicated that PAR2, through PI3K and AKT, lead to release of MMPs, which in turn lead to maturation of HB-EGF (Chung, Ramachandran, Hollenberg, & Muruve, 2013; Rattenholl et al., 2007), although an MMP-independent mechanism through Src has also been proposed (van der Merwe, Hollenberg, & MacNaughton, 2008). Our model for how Matriptase invokes cellular responses is highly likely to be incomplete. Indeed, others have indicated EGFR and AKT are downstream of Matriptase function (List et al., 2005; Schepis et al., 2018). We do not think these conflict with our model but will interface with it. A number of reports have demonstrated that PI3K/AKT and MEK/ERK function in parallel downstream of PAR2 (Sheng et al., 2019; Tanaka, Sekiguchi, Hong, & Kawabata, 2008; van der Merwe, Moreau, & MacNaughton, 2009). Furthermore, there is evidence that PKC activates both MEK/ERK and EGFR independently following PAR2 stimulation (Al-Ani et al., 2010), and that PI3K is activated by PAR2 via Gq (P. Wang & DeFea, 2006). Cell identity, sub-cellular localisation, β-arrestin scaffolding and biased agonism/antagonism are known to generate alternative downstream outputs from PAR2 (Zhao et al., 2014). To understand fully the roles of Matriptase and PAR2 in epithelial homeostasis and carcinoma, it will be critical to map how, when, and where they activate different downstream pathways.

**Figure 11:**
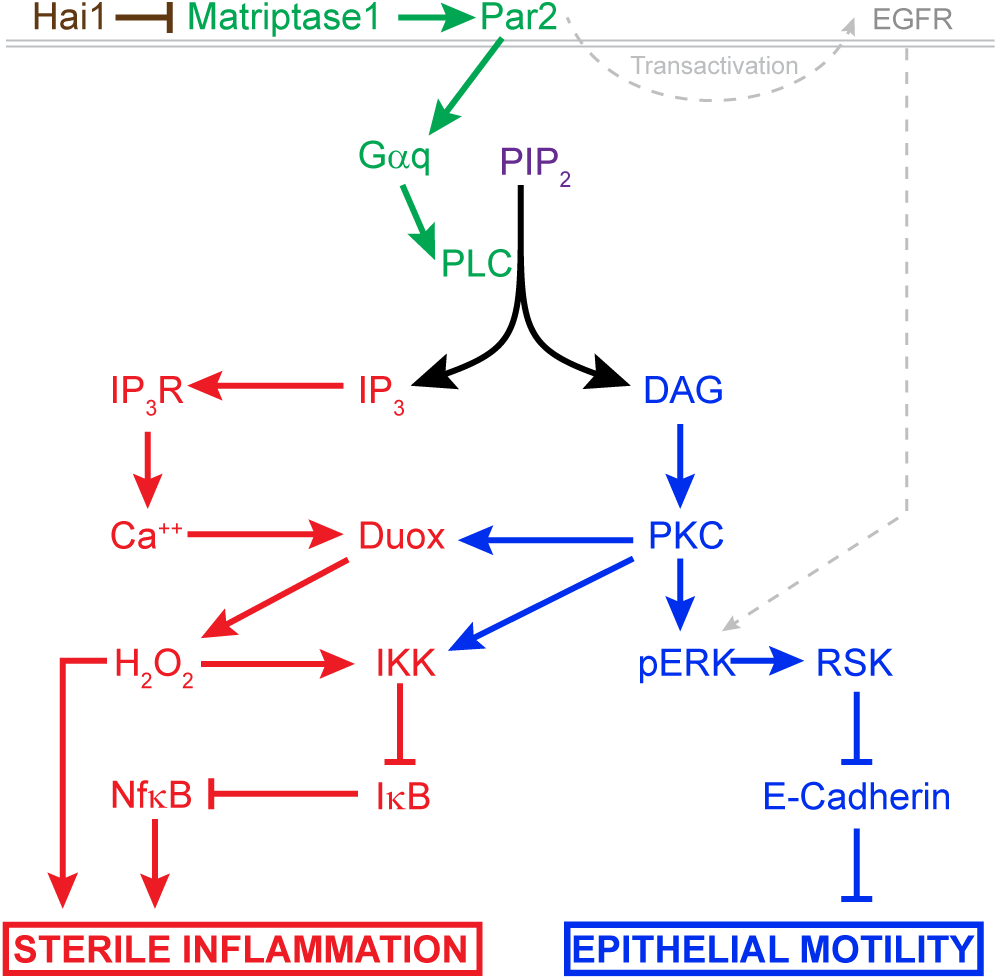
Model of pathway activated downstream of Hai1 and Matriptase. Proposed model of pathways downstream of Hai1 which drives Sterile Inflammation (red) and Epithelial Motility (blue). A previously defined transactivation of EGFR is also integrated. Other pathways known to act downstream of Matriptase, involving cMet, PI3K, AKT and mTOR are not shown.

## Materials and Methods

### Zebrafish husbandry and lines

Fish were housed at the IMCB and the NTU zebrafish facilities under IACUC numbers #140924 and #A18002 respectively, and according to the guidelines of the National Advisory Committee for Laboratory Animal Research. Embryos were derived by natural crosses and staged as per Kimmel et al (Kimmel, Ballard, Kimmel, Ullmann, & Schilling, 1995) and raised in 0.5x E2 medium (7.5mM NaCl, 0.25mM KCl, 0.5mM MgSO_4_, 75μM KH2PO4, 25μM Na_2_HPO_4_, 0.5M CaCl_2_, 0.35mM NaHCO_3_). Anaesthesia was administered in E2 medium (embryos) or fish tank water (adults) using 0.02% pH7.0 buffered Tricaine MS-222 (Sigma). The *hai1a/ddf* alleles used were *hai1a*^*hi2217*^, *hai1a*^*fr26*^, *ddf*^*ti257*^, and *ddf*^*t419*^. For imaging neutrophils and keratino-cytes, the transgenic lines *Tg(mpx:EGFP)*^*i114*^ (Renshaw et al., 2006) and *Tg(krtt1c19e:lyn-tdtomato)*^*sq16*^ (Lee et al., 2014) were used, whilst early leukocytes were imaged with *Tg(fli1:EGFP)*^*y1*^ (Redd, Kelly, Dunn, Way, & Martin, 2006). To image NfkB pathway activity, the *Tg(6xHsa.NFKB:EGFP)*^*nc1*^ sensor line was used (Kanther et al., 2011).

### Genomic DNA and RNA extraction, Reverse transcription, and PCR

Adult fin clips or embryos were isolated following anaesthesia, and genomic DNA extracted by incubation at 55°C for 4hrs in Lysis buffer (10mM Tris pH8.3, 50mM KCl, 0.3% Tween20, 0.3% Nonidet P-40, 0.5μg/μl Proteinase K). PCRs were performed using GoTaq (Promega) on a Veriti thermal cycler (Applied Biosystems), and purified with a PCR purification kit (Zymo Research). TRIzol™ (Invitrogen) was used for RNA extraction following provided protocol, and cDNA generated from 1μg total RNA using SuperScript III Reverse Transcriptase (Invitrogen) with random hexamer primers. For qPCR, iTaq SYBR green (Bio-Rad) was used to amplify, with reaction dynamics measured on a Bio-rad CFX96 Real-Time PCR Detection System. For measuring *nfkbiaa* mRNA by qPCR, the following primers (5’ to 3’) were used to amplify a region encoded on exons 4 and 5: F-AGACGCAAAGGAGCAGTGTAG, R-TGTGTGTCTGCCGAAGGTC. Reference gene was *eef1a1l1* and the primers used amplified between exon 3 to 4: F-CTGGAGGCCAGCTCAAACAT, R-AT-CAAGAAGAGTAGTACCGCTAGCATTAC.

### RNA synthesis

RNA for *GCaMP6s* was synthesised from a pCS2 plasmid containing the *GCaMP6s* coding sequence (gift of the Solnica-Krezel Lab; (Chen, Xia, Bruchas, & Solnica-Krezel, 2017)). This was linearised with NotI (NEB), RNA *in vitro* transcribed with mMESSAGE mMACHINE SP6 Transcription Kit (Ambion) and purified by lithium chloride precipitation.

### Embryo injection and Morpholino

Embryos were aligned on an agarose plate and injected at the 1-cell stage with RNA or Morpholino diluted in Phenol Red and Danieau’s buffer using a PLI-100 microinjector (Harvard Apparatus). Injection needles were pulled from borosilicate glass capillaries (0.5mm inner diameter, Sutter) on a Sutter P-97 micro-pipette puller. The Duox morpholino (AGTGAATTAGAGAAATGCACCTTTT) was purchased from GeneTools and injected at 0.4mM with 0.2mM of the tp53 morpholino (GCGCCATTGCTTTGCAAGAATTG).

### Zinc Finger and TALEN Mutagenesis

To generate the *st14a* (=*matriptase1a)* mutant, CompoZr Knockout Zinc Finger Nucleases were designed and manufactured by Sigma-Aldrich, targeting the following sequence in Exon 6: CAGTTCCAGCAGCA-CACGaagcaGCAGTGGATCAGGCTGTG, where the forward and reverse strand ZFN binding sites are in uppercase and the cut site is in lowercase. To generate the *ikbkg* mutant, TALEN vectors targeting the sequence ATGGAGGGCTGG in second exon were designed and constructed by ToolGen (http://toolgen.com). TALEN vectors were linearized with *PvuII* (NEB) and purified using a PCR purification kit (Zymo Research), and then used for *in vitro* transcription with the MEGAshortscript T7 kit (Ambion). About 170-300pg of supplied ZFN RNAs or purified TALEN RNAs were then injected into one-cell stage WT zebrafish embryos, which were raised to 24hrs, then genomic DNA extracted.

For detection of fish with edited loci, PCR was performed on genomic DNA of injected fish with primers flanking the target site, cloned by TA cloning into pGEMT-Easy (Promega) or pCR2.1-TOPO-TA (Invitrogen) and individual clones sequenced to establish efficiency. Other embryos were raised to adulthood and their offspring were similarly genotyped to identify founder mutants.

### Small molecule treatment

All compounds for treating embryos were dissolved in DMSO, diluted in 0.5x E2 Embryo Medium and embryos treated by immersion. The compounds, and concentrations used, with catalog numbers were: Diphenyleneiodonium Chloride (DPI), 40μM (D2926, Sigma); Thapsigargin, 6.25**μ** M (T9033, Sigma); Bisindolylmaleimide I (GF109203X), 85**μ** M (S7208, Selleckchem;); YM-254890, 32μM (10-1590-0100, Focus Bio-molecule); 2-Aminoethyl diphenylborinate (2-APB), 2.5μM (D9754, Sigma), BI-D1870, 1.2μM (Axon-1528, Axon Medchem); Dimethyl fumarate, 9μM (242926, Sigma); Phorbol 12-myristate 13-acetate (PMA), 37.5 or 125ng/ml (P8139, Sigma); U0126, 100μM (9903, Cell Signaling Technology); PD184352 (CI-1040), 1.3μM (S1020, Selleckchem). Unless otherwise stated, controls for all experiments were exposed to 0.5% DMSO carrier in 0.5x E2 Embryo Medium.

### Proteomic Analysis

Batches of 100 WT, *ddf*^*t419*^ and *ddf*^*ti257*^ embryos were collected at 24hrs and 48hrs, dechorionated, deyolked and protein extracted as per (Alli Shaik et al., 2014). Protein was precipitated in 100% methanol at 4°C, then resuspended in 2-D cell lysis buffer (30 mM Tris-HCl, pH 8.8, containing 7 M urea, 2 M thiourea and 4% CHAPS). 2-D DIGE and Mass spectrometry protein identification was performed by Applied Biomics (Hayward, CA). Protein samples were labelled with either Cy2, Cy3 or Cy5, mixed and then subjected to 2-D DIGE to separate individual proteins. Gels were scanned using Typhoon TRIO (Amersham BioSciences) and analysed by Image QuantTL and DeCyder (ver. 6.5) software (GE-Healthcare). Spots with more than 1.5 fold change were picked, in-gel trypsin digested and protein identification performed by MALDI-TOF mass spectrometry and MASCOT search engine in the GPS Explorer software (Matrix science).

### In situ Hybridisation

A probe corresponding to the final 1078 bp of *rps6ka3a* (*RSK2a;* NM_212786.1) was generated by cloning a PCR-derived cDNA fragment into in pGEMT-Easy (Promega), linearising with *ApaI* (NEB) and transcribing a DIG probe with SP6 RNA polymerase (Roche). Whole mount in-situ hybridisation developed with NBT/ BCIP (Roche) was performed as described (Thisse & Thisse, 2008).

### Immunofluorescent, Dye staining and TUNEL

For antibody staining, embryos were fixed in 4% paraformaldehyde overnight at 4°C and then washed in PBT (0.1% Triton in PBS), permeabilized in −20°C Acetone for 7 minutes, washed in PBT, blocked for 3 hours in Block solution (PBT supplemented with 4% BSA and 1% DMSO), then incubated overnight at 4°C with primary antibody diluted in Block solution, washed extensively in PBT, re-blocked in Block solution then incubated overnight at 4°C with fluorescent secondary antibody diluted in Block solution. Following extensive PBT washing, embryos were cleared in 80% glycerol/PBS before imaging. Primary antibodies used and their dilutions are as follows: Chicken anti-eGFP antibody, 1:500 (ab13970, Abcam), Rabbit anti-eGFP, 1:500 (Tp401, Torrey Pines Biolabs), Rabbit anti-FITC, 1:200 (#71-1900, ThermoFisher), Rabbit anti-beta Catenin, 1:200 (ab6302, Abcam), Mouse anti-E-Cadherin, 1:200 (#610181, BD Biosciences), Mouse anti-Tp63, 1:200 (CM163, Biocare Medical), Rabbit anti-phospho-p44/42 MAPK (Erk1/2) (Thr^202^/Tyr^204^), 1:100 (#4370, Cell Signaling Technology), Rabbit anti-p44/42 MAPK (Erk1/2), 1:100 (#9102, Cell Signaling Technology), Rabbit anti-p90RSK (Phospho-Thr^348^), 1:100 (A00487, Genscript). All secondary antibodies were purchased from Invitrogen and used at 1:700 and were Alexa Fluor-488 Donkey anti-rabbit (A21206), Alexa Fluor-647 Donkey anti-rabbit (A31573), Alexa Fluor-546 Donkey anti-mouse (A10036) and Alexa Fluor-488 Goat anti-chicken (A-11039). Nuclei were counterstained using 5μg/ml of DAPI (4’,6-Diamidino-2-Phenylindole, Dihydrochloride; D1306, Invitrogen) added during secondary antibody incubation.

To stain hydrogen peroxide, embryos were incubated for 60min at room temperature with 12.5μM pentafluorobenzenesulfonyl fluorescein (PFBSF) (#10005983, Cayman Chemicals), then rinsed in Embryo Medium, anaesthetized and imaged.

Fluorescent TUNEL staining was performed using the Fluorescein In Situ Cell Death Detection Kit (11684795910, Roche), with the fluorescein detected by antibody staining using rabbit anti-FITC, and co-immunostained for TP63 and eGFP).

### Microscopy and Statistical analysis

Still and timelapse imaging was performed on upright Zeiss AxioImager M2, Zeiss Light-sheet Z.1, upright Zeiss LSM800 Confocal Microscope or Zeiss AxioZoom V16 microscopes. Embryos were mounted in 1.2% Low Melting Point Agarose (Mo Bio Laboratories) in 0.5x E2 medium in 35 mm glass-bottom imaging dishes (MatTek) or in a 1mm inner diameter capillary for Light-sheet timelapse. When imaging was performed on live embryos, the embryo media were supplemented with 0.02% Tricaine. Image processing was done using Zen 3.1 software (Zeiss), ImageJ (ver. 1.52p) or Imaris (Bitplane) and compiled using Photoshop CS6 (Adobe). Graphpad Prism was used for statistical analyses and graph generation. In all statistical tests, * = p<0.05, ** = p<0.01, *** = p<0.001.

## Supplementary Figures

**Supplementary Figure 1:**
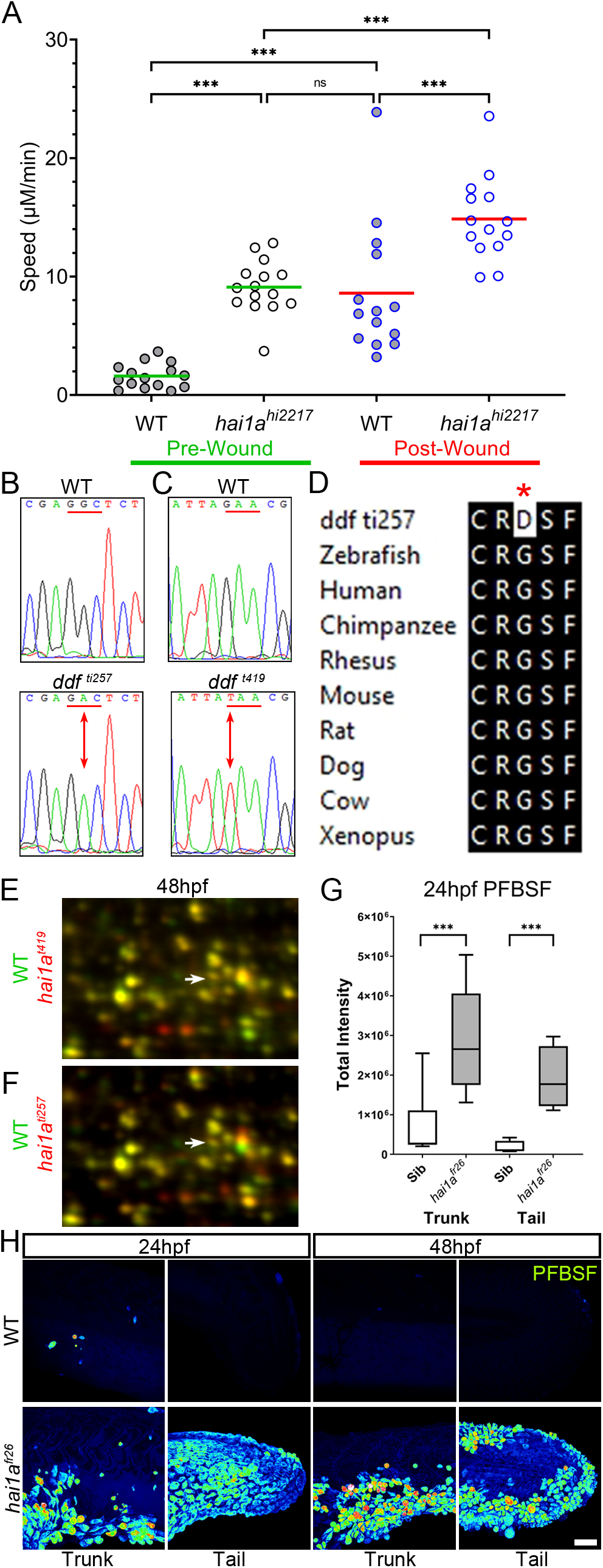
The epidermis of *hai1a* mutants displays elevated hydrogen peroxide. **A:** Graph of neutrophil speeds before and after wounding in WT and *hai1a*^*hi2217*^. Neutrophils in wounded WT move as fast as unwounded *hai1a* mutants. Wounding *hai1a* mutants accelerates neutrophils. **B-C:** Sequence chromatograms of genomic DNA from WT (upper panels) and *dandruff* alleles *ti257* (B) and *t419* (C) (lower panels). Altered codon is underlined in red with altered base indicated by arrow. **D:** ClustalW protein alignment of the amino acid substitution in the *ti257* allele showing broad conservation of across vertebrates. **E-F:** Selected regions of 2D gel of protein extracted from 48hpf embryos for *hai1a*^*t419*^ (E) or *hai1a*^*ti257*^ (F) in red, superimposed over WT protein samples (green in both). The shift in pI of Peroxiredoxin4 in both alleles is indicated with an arrow. **G:** Box and whiskers plot of PFBSF fluorescent staining intensity of WT and *hai1a*^*fr26*^ mutants at 24hpf in trunk and tail. n=9; t-test *** p<0.001. **H:** Lateral projected confocal views of PFBSF staining of WT (upper row) and *hai1a*^*fr26*^ (lower row) trunks and tails at 24hpf and 48hpf. Scale Bar H = 50μm.

**Supplementary Figure 2:**
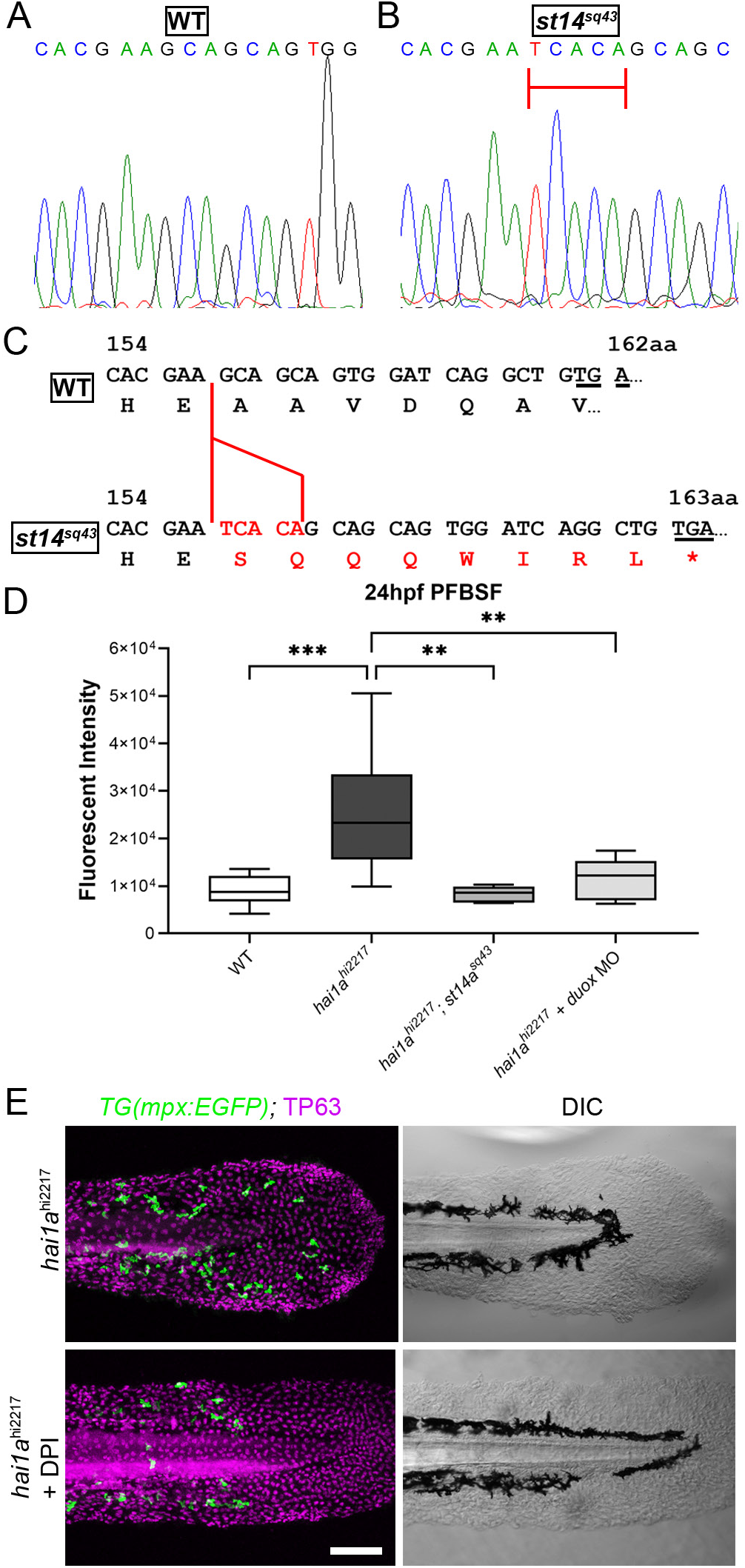
Generation of *st14a* mutant, and Duox inhibition reduces hydrogen peroxide and neutrophils in *hai1a* mutants. **A-B:** Sequence chromatograms of part of exon 6 of the zebrafish *st14a* gene from WT (A) and the *st14a*^*sq43*^ allele showing the 5bp insertion highlighted by red bar. **C:** Effect of the 5bp insertion (red) in the *sq43* allele leads to alteration of reading frame, 8 aberrant amino acids followed by a termination codon (red lettering). The codon altered to nonsense codon is underlined in both WT and mutant sequences. **D:** Box and whiskers plot of PFBSF fluorescent staining intensity of 24hpf WT, *hai1a*^*hi2217*^ mutants, *hai1a*^*hi2217*^; *st14a*^*sq43*^ double mutants and *hai1a*^*hi2217*^ injected with *duox* MO. n=8; ANOVA with Bonferroni post-test *** p<0.001, ** = p<0.01. **E:** TP63 (magenta) and eGFP (green) antibody staining (left column) with DIC imaging (right column) for 48hpf *hai1a*^*hi2217*^; *Tg(mpx:eGFP)*^*i114*^ mutants, either untreated (top row), or treated with 40μM DPI (bottom row). Scale bar D = 100μm.

**Supplementary Figure 3:**
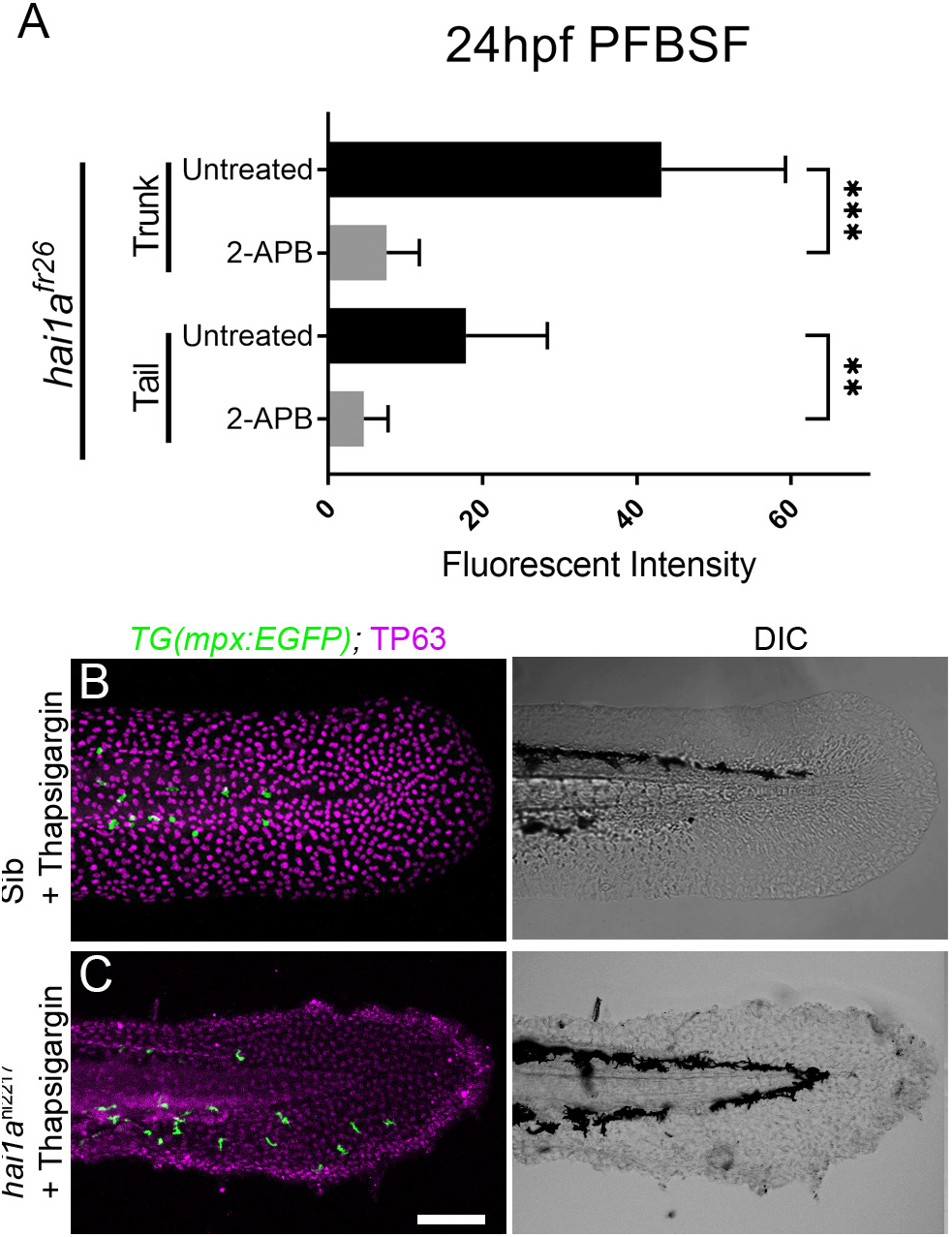
Thapsigargin and 2-APB reduces neutrophil inflammation but not epidermal defects in *hai1a* mutants. **A:** Plot of PFBSF fluorescent staining intensity of *hai1a*^*fr26*^ mutants at 24hpf in trunk and tail, either untreated or treated with 2-APB. n=14; ANOVA with Bonferroni post-test *** = p<0.001 ** = p<0.01. **B-C:** Projected confocal images of tail fins of 48hpf larvae immunostained with TP63 (magenta) and eGFP (green) (left column) with DIC imaging (right column). Larvae were both hemizygous for *Tg(mpx:EGFP)*^*i114*^ transgene and were treated with 6.5μM Thapsigargin. Genotypes are *hai1a*^*+/hi2217*^ sibling (A), *and hai1ahi^2217^* mutant (B). Rescue of neutrophil inflammation, but not epidermal defect is apparent in treated *hai1a* mutant. Scale bar B = 100μm.

**Supplementary Figure 4:**
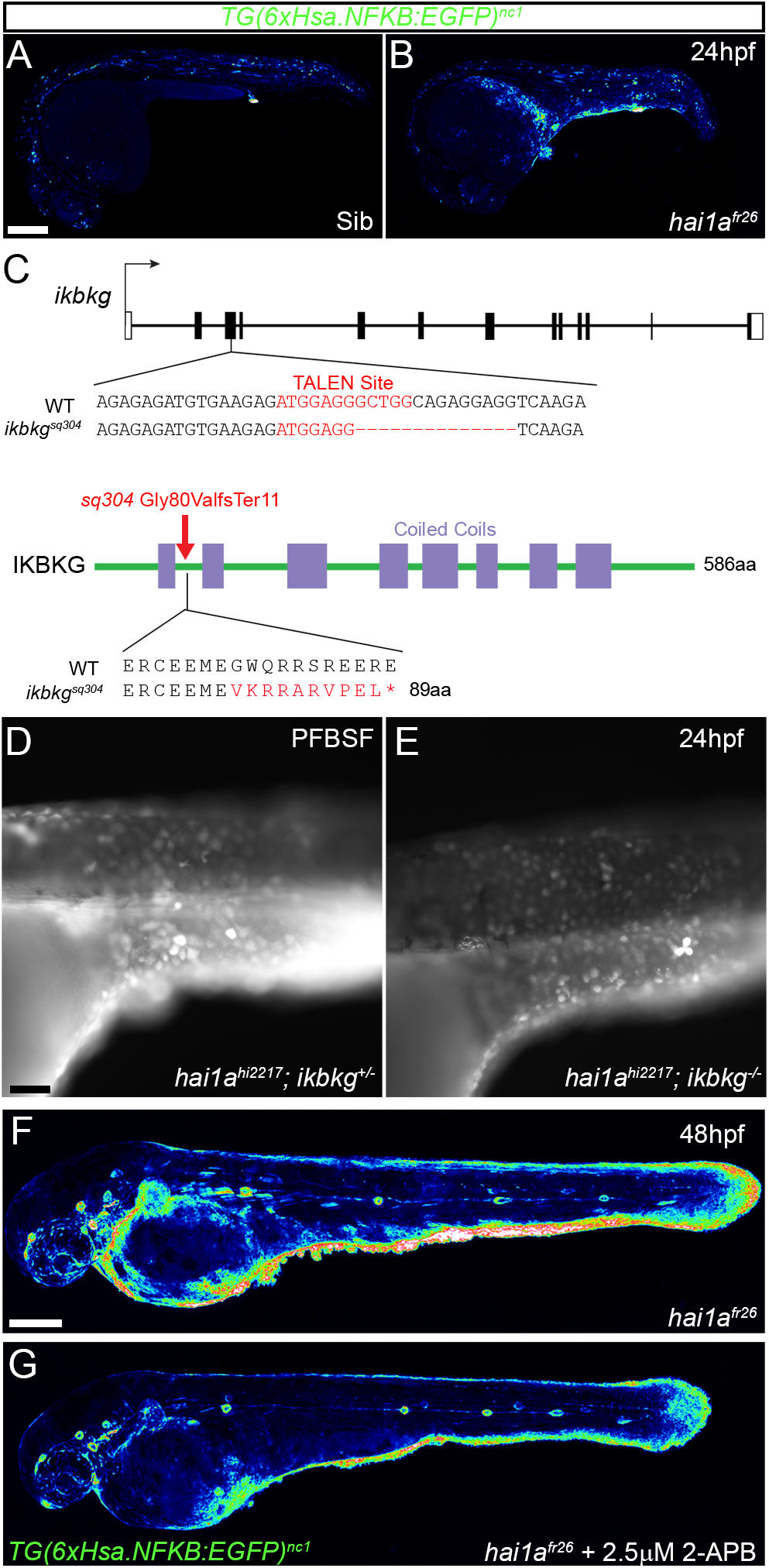
NfkB signalling is elevated in *hai1a* mutants and mutation of *ikbkg* rescues neutrophil inflammation. **A-B:** Lateral confocal projections of *Tg(6xHsa.NFKB:EGFP)*^*nc1*^ embryos reporting NfkB signalling levels of *hai1a*^*fr26*^ at 24hpf (B), and WT (A). **C:** Nature of the *ikbkg*^*sq304*^ mutant allele showing TALEN site location within the intron-exon structure of the gene (coding and non-coding exons depicted as filled and open boxes respectively). Sequence of part of exon 3 shown below with the TALEN binding site in red. The 14bp deletion in the *ikbkg*^*sq304*^ allele indicated under the WT sequence as dashes. This leads to a frameshift changing codon 80 from GGC (Gly) to GGT (Val), then 9 aberrant amino acids followed by a stop codon. **D-E:** Lateral widefield fluorescent images of *hai1a*^*hi2217*^; *ikbkg*^*+/sq304*^ (D) and *hai1a*^*hi2217*^; *ikbkg*^*sq304*^ (E) embryos at 24hpf stained with PFBSF showing no loss of H_2_O_2_ in *hai1a* mutants upon mutation of *ikbkg*. **F-G:** Lateral confocal projections of 48hpf *hai1a*^*fr26*^; *Tg(6xHsa.NFKB:EGFP)*^*nc1*^ embryos, treated with DMSO (F) or with 2.5μM 2-APB (G). Scale bars A, F = 200μm, D = 50μm.

**Supplementary Figure 5:**
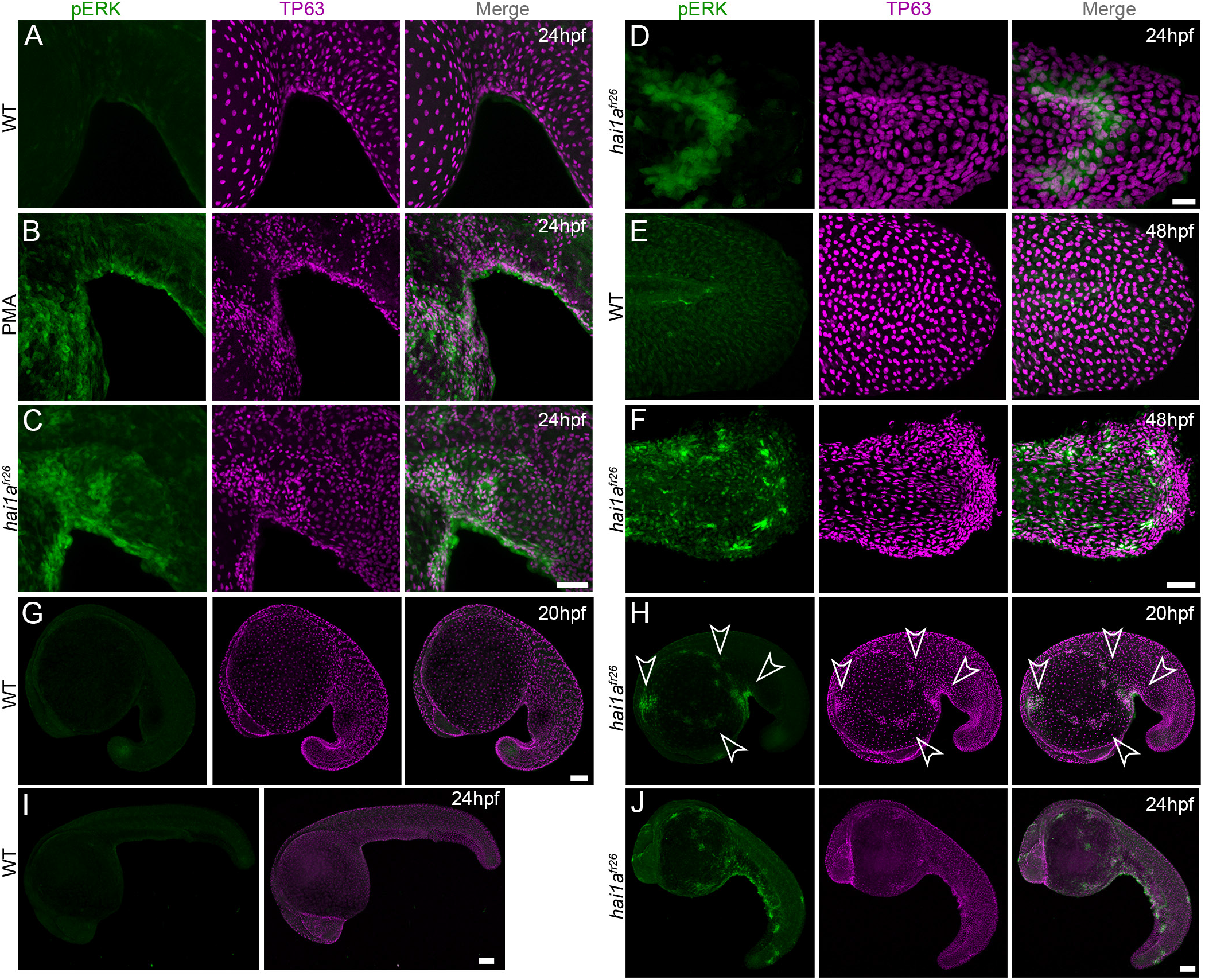
Elevation of pERK levels in PMA treated and *hai1a* mutant epidermis. **A-J:** Lateral projected confocal images of ventral trunks (A-C), tails (D-F) and whole embryos (G-J) immunostained for TP63 (magenta) and pERK (green) at 24hpf (A-D, I, J), 48hpf (E, F) and 20hpf (G, H). Both *hai1a*^*fr26*^ (C, D, F, H, J) and 125ng/ml PMA treated (B) embryos show increased epidermal pERK levels compared to untreated WT (A, E, G, I). Elevation of epidermal pERK is seen in *hai1a*^*fr26*^ mutants and PMA treated embryos as well as in nascent aggregates (arrowheads, H). Scale bars: C, F = 50μm; D = 20μm; G, I, J = 100μm.

**Supplementary Figure 6:**
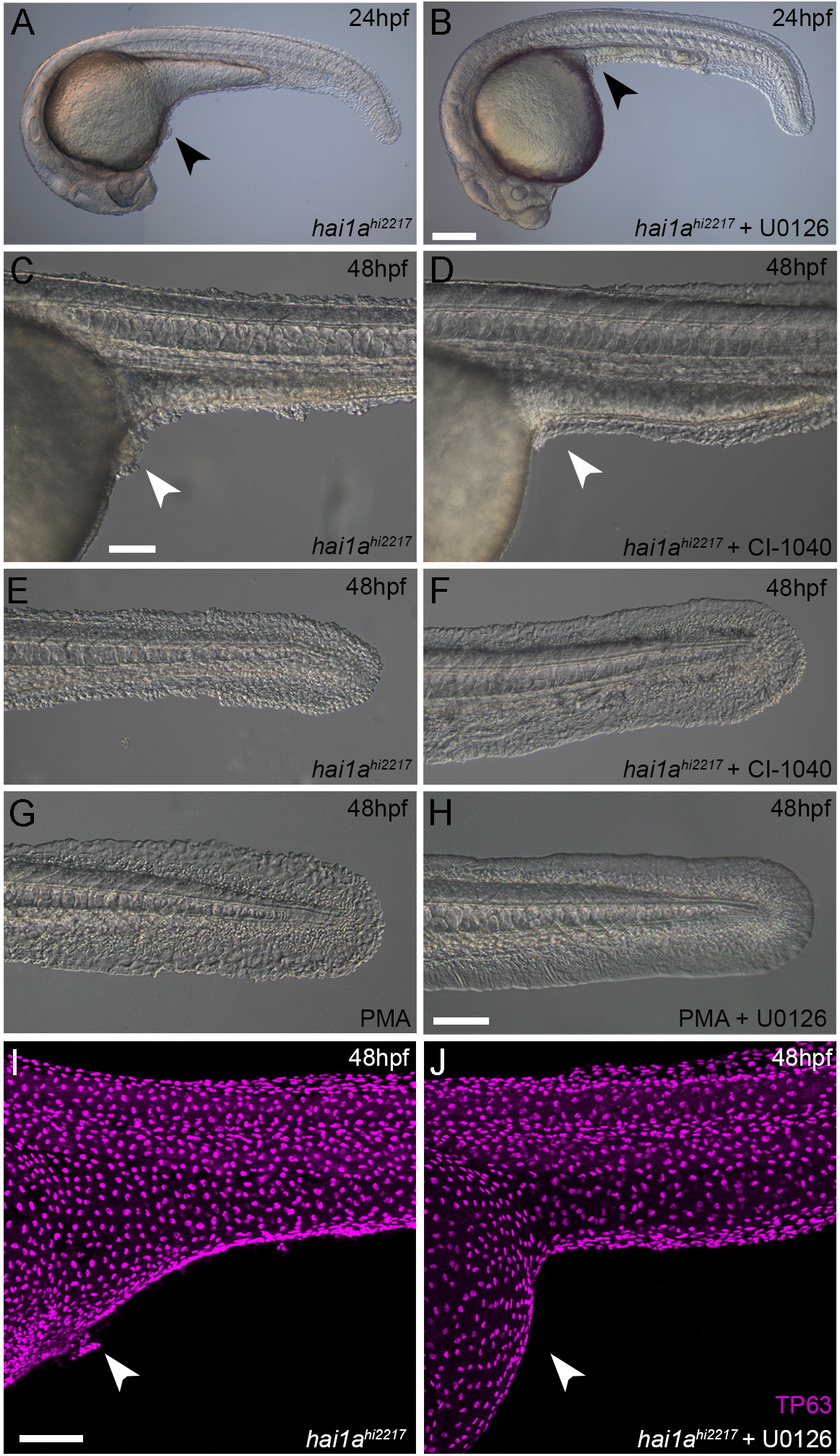
Rescue of the *hai1a* epidermal phenotype by pERK inhibitors. **A-F:** Lateral DIC images of 24hpf (A-B) or 48hpf (C-F) *hai1a*^*hi2217*^ embryos treated with either DMSO (A, C, E), U0126 (B) or CI-1040 (D, F) showing rescue of general morphology (B), trunk (D) and tail (F) epidermal phenotypes compared to DMSO treated *hai1a*^*hi2217*^. **G-H:** Lateral DIC images of tails of 48hpf WT treated with 125ng/ml PMA alone (G) or PMA and U0126 (H). **I-J:** Lateral projected confocal images of trunks of 48hpf *hai1a*^*hi2217*^ embryos treated with DMSO (I) or U0126 (J) and then fluorescently immunostained for TP63. Scale bars: B = 200μm; C, H, I = 100μm.

**Supplementary Figure 7:**
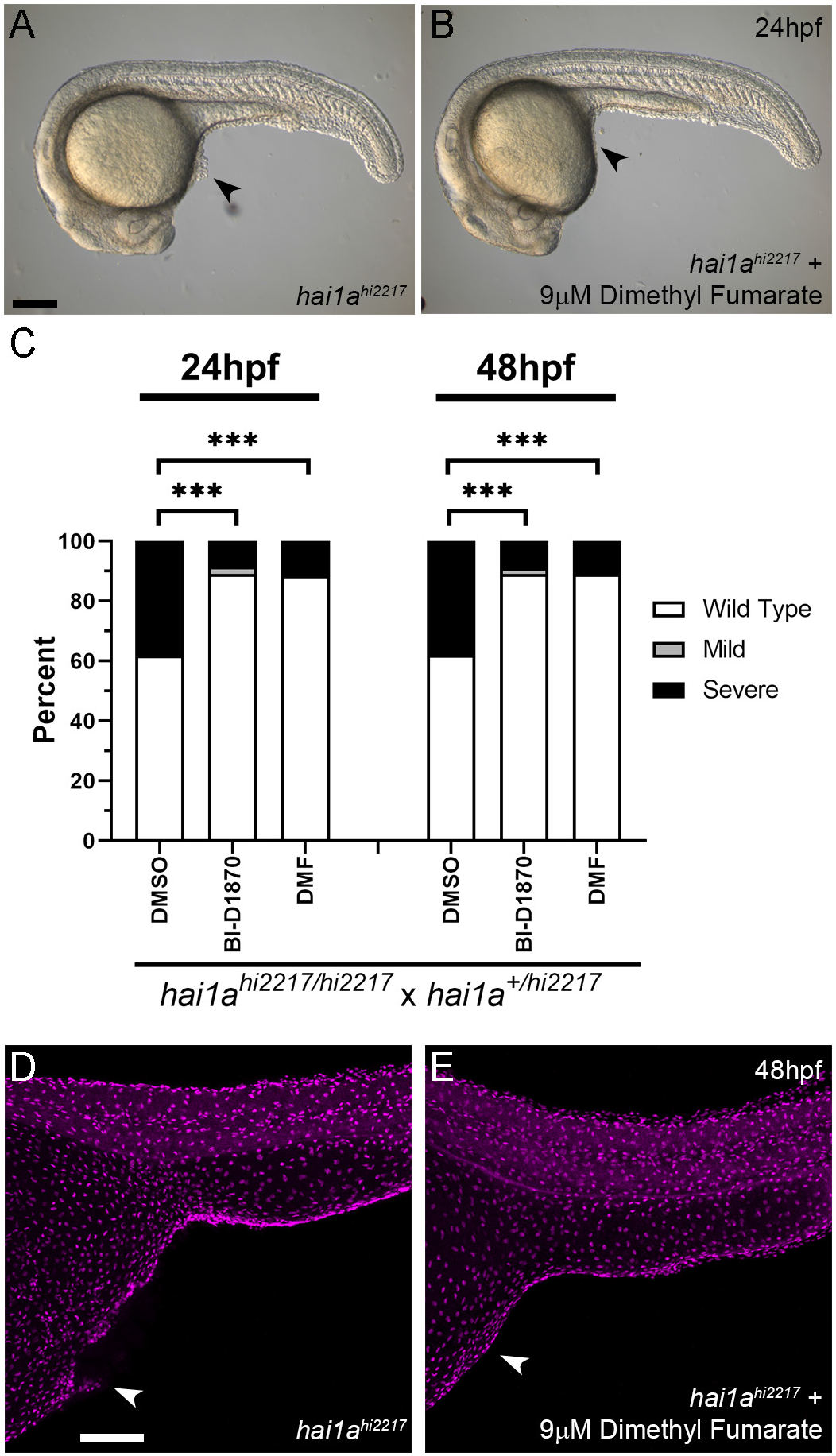
RSK inhibitors rescue the *hai1a* phenotype. **A-B:** Lateral DIC images of *hai1a*^*hi2217*^ embryos at 24hpf either untreated (A) or treated with 9μM Dimethyl Fumarate (B). Locations of epidermal aggregates and loss of tail fin morphology in *hai1a* mutants, and their rescue by RSK inhibitor treatment are indicated by arrowheads. **C:** Proportions of epidermal phenotypes from *hai1a*^*hi2217/hi2217*^ x *hai1a*^*+/hi2217*^ determined at 24hpf and 48hpf following treatments with DMSO, BI-D1870 or Dimethyl Fumarate (DMF). n=100; Chi-squared test; *** = p<0.001. **D-E:** Lateral projected confocal images of trunks of 48hpf *hai1a*^*hi2217*^ embryos, untreated (D) or treated with 9μM Dimethyl Fumarate (E), and then fluorescently immunostained for TP63. Scale bars: A=200μm; D = 100μm.

## Supplementary Table

**Supplementary Table1:** Prevalence of otolith and epithelial phenotypes in *hai1a* and *st14a* double mutants:

| *hai1a*<sup>+/hi2217</sup>;*st14a*<sup>+/sq43</sup> ♂ x *hai1a*<sup>hi2217/hi2217</sup>;*st14a*<sup>sq43/sq43</sup> ♀ |  |  |  |
| --- | --- | --- | --- |
| Observed (*Expected*) | WT Epidermis | *hai1a* epidermis | Total |
| Wildtype Otoliths | 72 (65) | 60 (65) | 132 (130) |
| No otoliths | 128 (65) | 0 (65) | 128 (130) |
| Total | 200 (130) | 60 (130) | 260 |

p<0.0001 (Chi -squared test)

## Supplementary Videos

**Supplementary Video 1: Neutrophils in WT and *hai1a*^*hi2217*^ 4dpf larva**

Projected confocal timelapses of eGFP positive neutrophils in the tail region of 4dpf *Tg(mpx:eGFP)*^*i114*^ (left) and *hai1a*^*hi2217*^; *Tg(mpx:eGFP)*^*i114*^ (right) larvae taken every 45 seconds for 45 minutes. Scale bar 50μm.

**Supplementary Video 2: Neutrophils in WT and *hai1a*^*hi2217*^ 4dpf larva before and after fin wound**

Projected confocal timelapses of eGFP positive neutrophils in the tail region of 4dpf *Tg(mpx:eGFP)*^*i114*^ (left) and *hai1a*^*hi2217*^; *Tg(mpx:eGFP)*^*i114*^ (right) larvae taken every 50 seconds for 250 minutes with the tail fin cut at 50 minutes. GFP is overlaid on DIC channel. Scale bar 50μm.

**Supplementary Video 3: Calcium dynamics in WT and *hai1a*^*fr26*^ embryos at 24hpf**

Projected confocal timelapses of eGFP in the trunks (left side) and tails (right side) of a 24hpf WT (top row) and *hai1a*^*fr26*^ (bottom row) embryos injected with *GCaMP6s* RNA, indicating calcium dynamics. Scale bar 50μm.

**Supplementary Video 4: Calcium dynamics in DMSO and 2-APB treated *hai1a*^*fr26*^ embryos at 24hpf**

Projected confocal timelapses of eGFP signal in the trunks (left side) and tails (right side) of 24hpf *hai1a*^*fr26*^ embryos injected with *GCaMP6s* RNA and treated with 0.03% DMSO (top row) and 2.5μM 2-APB (bottom row), indicating reduced calcium dynamics following 2-APB treatment. Scale bar 50μm.

**Supplementary Video 5: Basal keratinocyte membrane and neutrophil dynamics in 3dpf wild-type and *hai1a*^*hi2217*^ larvae carrying the *Tg(krtt1c19e:lyn-tdtomato)*^*sq16*^ and *Tg(mpx:eGFP)*^*i114*^ transgenes**

Projected light-sheet timelapses of the trunk of 3dpf WT (left) and *hai1a*^*hi2217*^ (right) larvae with neutrophils and basal keratinocyte membranes labelled by eGFP and Lyn-tdTomato respectively. Both larvae carried the *Tg(krtt1c19e:lyn-tdtomato)*^*sq16*^; *Tg(mpx:eGFP)*^*i114*^ transgenes. The *hai1a* mutants have highly dynamic neutrophils and keratinocyte membrane dynamics. Scale bar 20μm.

**Supplementary Video 6: Basal keratinocyte membranes in DMSO and PMA treated 3dpf *Tg(krtt1c19e:lyn-tdtomato)*^*sq16*^ larvae**

Zoomed projected light-sheet timelapses of basal keratinocyte membranes labelled by Lyn-tdTomato in the trunk of 3dpf *Tg(krtt1c19e:lyn-tdtomato)*^*sq16*^ larvae treated with 0.1% DMSO (left) and 37.5ng/ml PMA (middle and right) for 18hrs. Membranes are stable in DMSO treated larvae but were dynamic in PMA treated larvae. Images were captured every 20 seconds. Scale bar 10μm.

**Supplementary Video 7: Neutrophils and basal keratinocyte membranes in DMSO and PMA treated 3dpf *Tg(krtt1c19e:lyn-tdtomato)*^*sq16*^; *Tg(mpx:eGFP)*^*i114*^ larvae**

Lateral projection of light-sheet timelapse of neutrophils labelled by eGFP and basal keratinocyte cell membranes labelled by lyn-tdTomato in the trunks of 3dpf *Tg(krtt1c19e:lyn-tdtomato)*^*sq16*^ larva treated with 0.1% DMSO (left) and 37.5ng/ml PMA (right) for 18hrs. PMA treatment leads to slightly dynamic cell membranes and motile neutrophils. Images were captured every 20 seconds for 30 minutes. Scale bar 50μm.

